# Translation of non-canonical open reading frames as a cancer cell survival mechanism in childhood medulloblastoma

**DOI:** 10.1101/2023.05.04.539399

**Authors:** Damon A. Hofman, Jorge Ruiz-Orera, Ian Yannuzzi, Rakesh Murugesan, Adam Brown, Karl R. Clauser, Alexandra L. Condurat, Jip T. van Dinter, Sem A.G. Engels, Amy Goodale, Jasper van der Lugt, Tanaz Abid, Li Wang, Kevin N. Zhou, Jayne Vogelzang, Keith L. Ligon, Timothy N. Phoenix, Jennifer A. Roth, David E. Root, Norbert Hubner, Todd R. Golub, Pratiti Bandopadhayay, Sebastiaan van Heesch, John R. Prensner

**Author notes:** Address correspondence to: John R. Prensner, MD, PhD Department of Pediatrics, Medical Science Research Building II, Room 2560B 1150 Medical Center Drive, Ann Arbor, MI 48109, Sebastiaan van Heesch, PhD, Princess Máxima Center for Pediatric Oncology Heidelberglaan 25, 3584 CS Utrecht The Netherlands. These authors contributed equally (Damon A. Hofman, Jorge Ruiz-Orera).

## Abstract

A hallmark of high-risk childhood medulloblastoma is the dysregulation of RNA translation. Currently, it is unknown whether medulloblastoma dysregulates the translation of putatively oncogenic non-canonical open reading frames. To address this question, we performed ribosome profiling of 32 medulloblastoma tissues and cell lines and observed widespread non-canonical ORF translation. We then developed a step-wise approach to employ multiple CRISPR-Cas9 screens to elucidate functional non-canonical ORFs implicated in medulloblastoma cell survival. We determined that multiple lncRNA-ORFs and upstream open reading frames (uORFs) exhibited selective functionality independent of the main coding sequence. One of these, ASNSD1-uORF or ASDURF, was upregulated, associated with the MYC family oncogenes, and was required for medulloblastoma cell survival through engagement with the prefoldin-like chaperone complex. Our findings underscore the fundamental importance of non-canonical ORF translation in medulloblastoma and provide a rationale to include these ORFs in future cancer genomics studies seeking to define new cancer targets.

**Highlights:**

- Ribo-seq reveals widespread translation of non-canonical ORFs in medulloblastoma
- High-resolution CRISPR tiling reveals uORF functions in medulloblastoma
- ASNSD1-uORF controls downstream pathways with the prefoldin-like complex
- ASNSD1-uORF is necessary for medulloblastoma cell survival

## Introduction

High-risk medulloblastoma remains one of the most recalcitrant pediatric cancers, and children with MYC-amplified disease frequently succumb to relapsed disease.^1–3^ Besides MYC amplification, in-depth analyses of the medulloblastoma coding genome have identified and characterized additional somatic events in subsets of patients. Still, most tumors lack targetable mutations and do not yield insights regarding their aggressive behavior.^4–6^ At the same time, medulloblastoma is known to exhibit extensive rewiring of RNA translational control both through genetic mutation of the DDX3X RNA helicase in the WNT and SHH subtypes,^6–8^ as well as in Group 3/4 tumors through activation of the MYCN or MYC transcription factors, where recent genetic evidence indicates that control of RNA translation may be the most critical aspect of MYC function during tumorigenesis.^9–11^ This deregulation of RNA translational control in medulloblastoma leads not only to a wide discrepancy between RNA and proteomic signatures,^12, 13^ but also to a distinctive reliance on RNA translation factors^14^ and potential therapeutic options.^15, 16^

While translation of known proteins has been the focal point for prior research in medulloblastoma as well as other childhood brain cancers, the human genome also contains thousands of non-canonical open reading frames (ORFs).^17^ These previously understudied ORFs are ubiquitous regions of ribosome translation that occur separately from the known protein-coding sequences and have the capacity to influence gene activity or to encode proteins with distinct biological functions.^18–21^ For example, individual cancer-associated ORFs may generate novel cancer targets that influence cell phenotypes,^22, 23^ whereas other classes of ORFs are critical effectors of oncogene-induced gene regulation.^24^ However, the overall potential impact of such ORFs across and within cancers has not been determined.

Here, we have investigated the functional impact of translation of non-canonical ORFs in medulloblastoma. We demonstrate that these ORFs are commonly translated in medulloblastoma model systems and patient tumors, with translational control influenced by disease subtype. Using genome-wide CRISPR screens and ORF-specific saturation mutagenesis with CRISPR, we found that non-canonical ORFs are frequently essential for cell survival in medulloblastoma and describe widespread reliance on upstream open reading frames (uORFs) in particular. From these, we identify a uORF in the *ASNSD1* gene that is selectively upregulated and required for maintenance of cell survival by coordinating the function of the prefoldin-like complex, a poorly understood complex implicated in post-translational control.^25–27^ Together, our findings demonstrate that oncogenic uORFs can act as critical disease mediators both in medulloblastoma and, by extension, human cancers more broadly.

## Results

### Comprehensive translational profiling of medulloblastoma highlights biological subtypes

To characterize signatures of RNA transcription and translation in medulloblastoma, we profiled 32 unique patients/cell lines (14 medulloblastoma cell lines and 18 tumor samples) using RNA-seq and ribosome profiling^28^ (Figure 1A and **Table S1A-F**). Samples reflected major histological and molecular subtypes, including large cell/anaplastic and desmoplastic nodular, and MYC amplified subtypes (**Table S1A**). In total, we sequenced and mapped over 1.3 billion ribosome footprints across 32 samples (**Table S1A** and Figure S1A-C). For this, we further optimized the Ribo-seq procedure to capture high-quality ribosome footprints from low-input tumor samples down to 3 mg per sample (range: 3 - 75 mg). Ribosome profiling achieved an average of 78.8% in-frame reads (range 64.7 - 84.8%) with an average of 12,340 translated known protein-coding sequences (CDSs) quantified per sample (range 10,712 - 13,868 CDSs) (Figure 1B-D and Figure S1D-E). Tissue samples and cell lines exhibited similar performance metrics, with tumor samples yielding a higher number and thus greater diversity of detected CDSs (Figure 1C).

**Figure 1:**
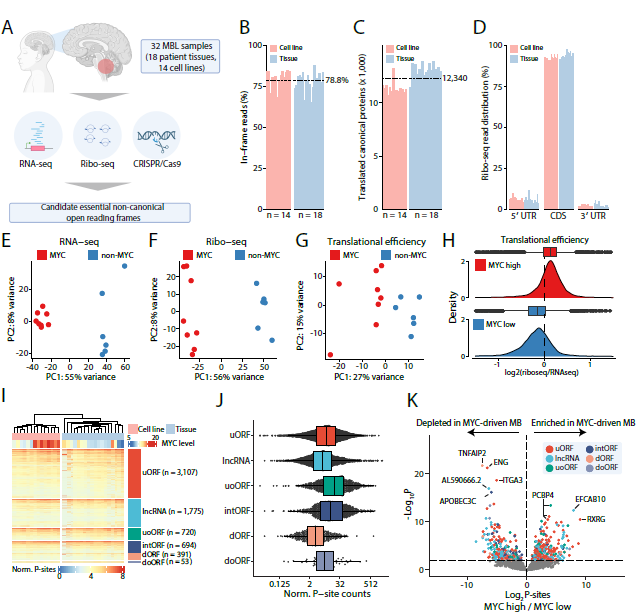
Comprehensive profiling of non-canonical ORF translation in medulloblastoma. A. Schematic depiction of experimental approach B. Bar plot showing the percentage of in-frame ribo-seq reads across all 14 cell line samples and 17 tissue samples. C. Bar plot showing the number of translated canonical proteins (defined as P-sites per million > 1) across all samples. D. Bar plots showing percentages of reads mapping to coding sequences (CDS) and untranslated regions (5’ UTR and 3’ UTR) of protein coding sequences across all samples. E. A principal component analysis (PCA) showing MYC-driven and nonMYC-driven samples using RNA-seq data. F. A PCA showing MYC-driven from nonMYC-driven samples using Ribo-seq data. G. A PCA separating MYC-driven from nonMYC-driven samples using translation efficiency values. Each dot represents one sample. H. A density plot showing the distribution of translational efficiency values for each gene in MYC-driven and non-MYC driven medulloblastoma subgroups. Boxplots show lower quartile, median, and upper quartile values, with whiskers extending to highest and lowest observations. I. Heatmap showing translation levels of translated non-canonical ORFs (rows) across all samples (columns). Rows and columns were clustered in an unsupervised manner within sample type (tissue and cell line) and ORF biotype groups. Samples are annotated by MYC translation levels. Translation levels are calculated as transformed normalized P-site counts. J. Boxplots showing distributions of translation levels of translated non-canonical ORFs, separated by ORF biotype. Each dot represents the mean translation level of one ORF across all samples. Boxplots show lower quartile, median, and upper quartile translation levels for each ORF biotype. Translation levels are calculated as normalized P-site counts. X-axis reflects a log2 scale. K. Volcano plot of changes in translation levels between MYC-driven and non-MYC driven medulloblastoma cell lines. Each dot reflects a single non-canonical ORF, colored by ORF biotype. Dots above the dashed horizontal line have an FDR < 0.01. Labels for top 5 upregulated (log2 fold change > 2) ORFs with lowest p_adj_ and top 5 downregulated ORFs (log2 fold change < −2) with lowest p_adj_ are shown. **See also** Figure S1.

Clustering of cell lines by mRNA expression levels as well as ribosome profiling demonstrated distinct biological signatures between MYC-driven and non-MYC-driven cell lines (Figure 1E-F). Given prior proteogenomic data demonstrating discrepant RNA and protein signatures in medulloblastoma,^12, 13^ we next determined mRNA translational efficiency scores by comparing ribosome profiling and RNA-seq data (see **Methods** and **Table S1G**), and observed clustering of MYC-driven compared to non-MYC cell lines, indicative of stark differences in translational control between medulloblastoma subtypes driven by MYC activity (Figure 1G). Indeed, compared to non-MYC-driven cells, MYC-driven cell lines exhibited a significantly increased mRNA translational efficiency overall (Figure 1H; Wilcoxon test; p < 2.2 × 10^−16^). Consistent with these results, Gene Ontology and Gene Set Enrichment Analyses highlighted pathways related to ribosome biogenesis, translation initiation and elongation, and neuronal differentiation as distinctive between subtypes depending on MYC activity (Figure S1F, **Table S1H**). Together, these data support prior observations that dysregulated RNA translational control is widespread in medulloblastoma and reflects underlying differences in tumor subtype biology.^12, 13^

### Translation of non-canonical ORFs is common in medulloblastoma

Motivated by increasing reports of functional non-canonical ORFs detected through translational profiling,^21, 22, 29, 30^ we next sought to quantify the contribution of these ORFs to the medulloblastoma translatome. We assessed translation of 8,008 non-canonical ORFs derived from our previous analyses^22^ as well as a recently compiled human consensus ORF^17^ catalog using our tissue and cell line ribosome profiling datasets. We observed translation for 7,530 non-canonical ORFs in at least 1 sample and 6,740 in at least 5 samples (Figure 1I-J; **Table S1I-J**). Among these, translation of uORFs was most commonly and reproducibly detected (n = 3,107; ≥5 samples), followed by translation of lncRNA ORFs (n = 1,775), upstream overlapping ORFs (uoORFs, n = 720), internal ORFs (intORFs, n = 694), downstream ORFs (dORFs, n = 391), and downstream overlapping ORFs (doORFs, n = 53). Importantly, translational efficiency analysis of non-canonical ORFs recapitulated disease clusters, similar to annotated CDSs, indicating subtype-specific control of non-canonical ORF translation (Figure S1G). Overall, 717 non-canonical ORFs displayed differential translation levels between subtypes (p_adj_ < 0.01), with 268 ORFs showing increased translation in MYC amplified medulloblastoma (p_adj_ < 0.01, log2 fold change > 2) (Figure 1K and **Table S1K-L**). This indicates that the medulloblastoma translatome is populated by thousands of diverse non-canonical ORFs and that translation of non-canonical ORFs is a characteristic feature of medulloblastoma disease subtypes.

### Non-canonical ORFs are essential and specific in medulloblastoma cell survival

Non-canonical ORFs are increasingly recognized as serving key roles in cancer cell biology, in some cases through the generation of a stable bioactive protein.^19, 22, 23, 31^ Given their frequent and subtype-specific translation in medulloblastoma, we next sought to nominate non-canonical ORFs with key functional roles in this disease. We designed a genome-wide CRISPR guide RNA library targeting 2,019 ORFs and conducted loss-of-function knockout screening in 7 medulloblastoma cell lines (4 MYC-driven, 3 non-MYC-driven) in order to nominate non-canonical ORFs implicated in medulloblastoma cancer cell survival (Figure 2A, Figures S2A-C, and **Table S2A**). Performance metrics of the CRISPR screens were similar across cell lines and demonstrated high biological reproducibility (Figures S2D-J and **Tables S2B-E**).

**Figure 2:**
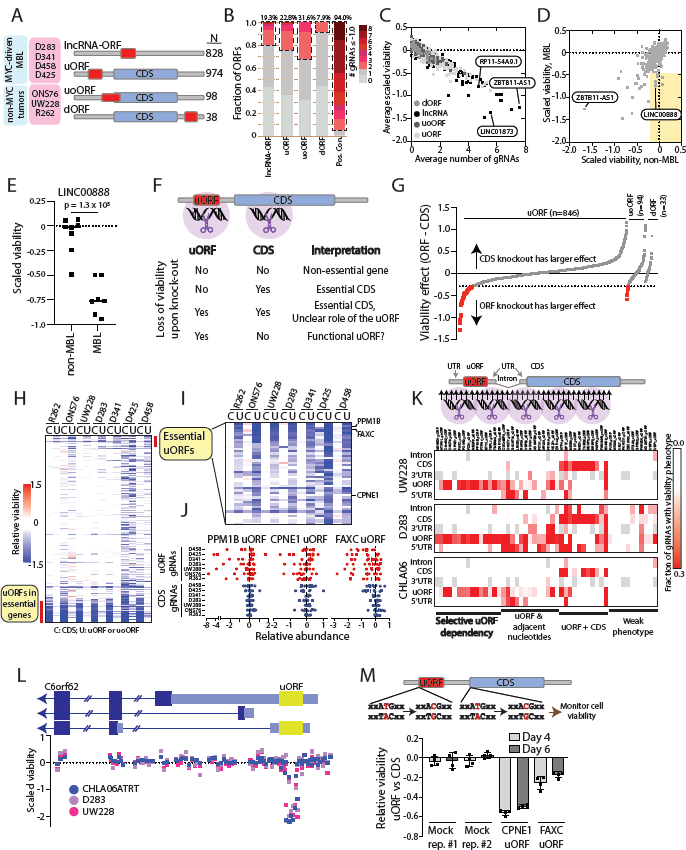
Non-canonical ORFs are frequently essential genes in medulloblastoma. A. A schematic description of the cell lines and numbers of non-canonical ORFs evaluated by CRISPR screening. B. A bar plot showing frequency of essentiality among different classes of non-canonical ORFs. At least 2 gRNAs had to score as depleted to nominate an essential non-canonical ORF. C. A scatter plot showing the relationship between the average ORF knockout phenotype across cell lines compared to the average number of gRNAs with a viability score of <= −0.5 across cell lines. Previously identified ORFs from^22^ are indicated. D. A scatter plot showing the correlation of ORF knockout phenotypes across a previously published panel of 8 non-medulloblastoma cancer cell lines^22^ and the current dataset of 7 medulloblastoma cell lines. Medulloblastoma-specific effects are highlighted in the yellow box. E. The impact of knockout of an ORF in LINC00888 in medulloblastoma and non-medulloblastoma cancer cell lines.^22^ Each dot reflects an individual cell line. The Y axis reflects the overall loss-of-viability phenotype of LINC00888 knockout. P value by a two-tailed Student’s T-test. F. A schematic reflecting the knockout strategy to identify uORFs and uoORFs with putative functional consequences in medulloblastoma cell viability. G. A line graph showing the scaled loss of viability when comparing knock-out of a uORF, uoORF, or dORF with knock-out of the associated parental coding sequence (CDS) for that gene. The Y axis shows the differential in viability effect. The X axis reflects each individual ORF. H. A heatmap showing scaled loss of viability for each pair of a parental CDS and a uORF or uoORF across all tested cell lines. Pan-essential CDSs are indicated. C, parental CDS; U, uORF or uoORF. I. An expanded view of the heatmap in (H), focusing on cases in which knock-out of a uORF or uoORF resulted in substantially more loss of viability compared to knock-out of the parental CDS. J. Individual gRNA level data for three essential uORFs. Here, each dot represents a gRNA to either the indicated uORF or the associated CDS. The Y axis shows the cell line for the data points. The X axis shows the scaled loss of viability associated with the gRNA. K. *Top*, a schematic showing the tiling saturation gRNA library design. *Bottom*, a heatmap showing the fraction of gRNAs for the given genomic region of the indicated ORF that scored as displaying a loss of viability phenotype. ORFs are organized along the X axis according to whether they exhibited a selective knock-out phenotype, a phenotype in conjunction with other gRNAs, or a weak phenotype. L. Individual gRNA-level data from the tiling saturation screen for the C6orf62 uORF. Each dot represents a gRNA. The Y axis shows the loss-of-viability associated with each gRNA. gRNAs are ordered along the X axis to align with the schematic of the C6orf62 gene and uORF. M. Base editing of the CPNE1 and FAXC uORF start codons or the start codons of their associated parental CDSs in D425 medulloblastoma cells. The barplot displays the differential in viability for uORF compared to CDS gRNA.

In aggregate, 390 ORFs (21.4%) demonstrated an essentiality phenotype in at least one cell line, with 112 out of 390 of ORFs displaying an effect on cell survival in at least 2 independent cell lines (Figure 2B and **Tables S2E-G**). Overall, upstream overlapping ORFs (uoORFs) and uORFs had higher rates of essentiality, although this observation was likely influenced by proximity to annotated CDSs and gene promoters (Figure S2K). dORFs, located in the 3′ UTRs of protein-coding mRNAs, exhibited the lowest rates of essentiality (Figure 2B), consistent with their generally lower translation rates (Figure 1J).

Across all cell lines, the strongest loss-of-function phenotypes were observed by the known pan-lethal effect of *ZBTB11-AS1*, which we previously characterized as an 88 amino acid microprotein, as well as several other pan-lethal lncRNA-ORFs in *LINC01873* and *RP11-54A9.1* (Figure 2C).^22^ A direct comparison of 528 ORFs screened in our current cohort of 7 medulloblastoma cell lines and our prior cohort of 8 non-medulloblastoma cell lines^22^ revealed 14 ORFs whose knockout had a significantly increased loss-of-viability phenotype in the medulloblastoma cohort (Figure 2D and **Table S2H**). Among these, we observed particularly pronounced medulloblastoma-specific viability effects for *LINC00888*, which encodes a microprotein whose translation is particularly elevated in MYC-driven medulloblastoma samples (Figure 2E-F and Figure S2L). Thus, medulloblastoma may possess a unique landscape of non-canonical ORF functions.

### Selective gene dependency for upstream open reading frames in medulloblastoma

While functionality of ORFs in some lncRNAs has been well-established,^22, 29, 30, 32^ we were intrigued to note the abundant uORFs with an essentiality phenotype upon knockout (Figure 2B). As most uORFs and uoORFs are conventionally thought to be regulatory sequences for adjacent canonical CDSs,^18, 33^ recent studies have indicated that some uORFs contain sequence variants^34, 35^ and encode protein products^36–40^ that contribute to disease and function independent of the canonical CDS encoded by the same gene. We therefore sought to determine whether any uORFs or uoORFs harbored a selective cancer dependency phenotype that might suggest unique biological relevance of the ORF. To do this, we performed matched knockout of the uORF or uoORF and knockout of the adjacent CDS in 964 cases (>90% with at least 7 gRNAs per ORF) and compared the knockout phenotypes (Figure 2F and **Table S2A**).

We observed that 69 (7.2%) of uORFs or uoORFs exhibited a substantial loss-of-viability phenotype upon knockout that was not recapitulated by knockout of the adjacent CDS (Figure 2G-J and **Table S2F**), of which 29/69 (42.0%) represented pan-lethal effects observed in at least 6 cell lines. To probe this observation further, we generated a custom tiling gRNA library that saturated 50 of the 69 mRNAs (median 79.5 gRNAs per gene, range 68 - 112) in which the uORF exhibited a lethality phenotype and performed loss-of-function screens in three cell lines (1 non-MYC MBL, 1 MYC MBL, and 1 atypical teratoid/rhabdoid tumor) (Figure S2M-N and **Table S2I-K**). In total, 15 uORFs exhibited a knockout phenotype only when uORF-targeting gRNAs were used, corroborating the above-mentioned effects at a high resolution and indicating precise selective dependency relative to the CDS (Figure 2K, Figures S2O-Q, and **Table S2L**), as exemplified by the C6orf62 uORF tiling knockout results (Figure 2L). Using two additional examples of uORFs located in the *CPNE1* and *FAXC* genes, we also verified that uORF translation was the critical feature for dependency through base editing of the uORF start codon (Figure 2M). These results indicate that a subset of uORFs may have unique roles in medulloblastoma cell viability.

### Identification of a uORF in ASNSD1 as a genetic dependency in medulloblastoma

By comparing MYC- and non-MYC driven cell lines, we were intrigued to observe that MYC-driven medulloblastoma cells exhibited enhanced essentiality phenotypes with uORF knockout (p < 0.001, Mann Whitney U test), but not for uoORFs or dORFs (Figure 3A). While most differential uORF essentiality phenotypes were modest in magnitude, we found 4 uORFs exhibiting a statistically-significant enrichment in MYC-driven cells (Figure 3B). Among these, a uORF in the *ASNSD1* gene exhibited particular strength as a vulnerability gene in MYC-driven medulloblastoma (Figure 3C). This gene also demonstrated among the most differential phenotypes between uORF knockout and main CDS knockout, with a highly selective phenotype (Figure 3D and Figure S3A).

**Figure 3:**
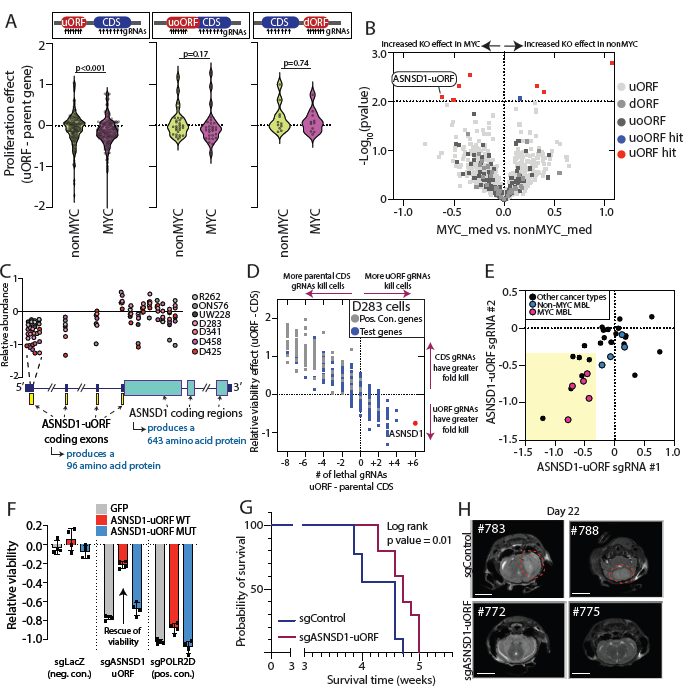
ASNSD1-uORF drives medulloblastoma cell survival. A. Violin plots showing the differential viability phenotype in MYC- or nonMYC-driven medulloblastoma cells for knock-out of uORFs, uoORFs, or dORFs that scored as hits in the CRISPR screen. P values by a Mann Whitney U test. B. A volcano plot showing the differential viability phenotype of knock-out of uORFs, uoORFs, and dORFs in MYC- and non-MYC cell lines. Hits are indicated with the shown colors. P values are by a two-tailed Student’s t-test. C. Individual gRNA-level data for ASNSD1-uORF and ASNSD1 parental CDS in the primary CRISPR screen. Each dot reflects a gRNA; dot colors reflect the indicated cell lines. The Y axis shows scaled viability after knock-out with each gRNA. The X axis reflects the genomic position of the gRNA relative to the ASNSD1 gene structure shown below. D. A scatter plot comparing the magnitude of viability phenotype of uORF knock-out relative to parental CDS knock-out in D283 cells. The X axis shows the number of gRNAs inducing a loss-of-viability phenotype for the uORF minus that number for the parental CDS. The Y axis shows the average loss-of-viability phenotype of the 4 most effective gRNAs for the uORF minus that number of the parental CDS. Positive control genes are shown in gray and other uORF genes are shown in blue. E. A scatter plot showing the degree of loss-of-viability for ASNSD1-uORF knock-out using two gRNAs across 33 cell lines. MYC-driven medulloblastoma cells are shown in pink and nonMYC medulloblastoma cells are shown in blue. Other cell lines are shown in black. F. A barplot showing the loss-of-viability for ASNSD1-uORF knock-out in D341 cells stably overexpressing GFP, ASNSD1-uORF, or AUG-mutant ASNSD1-uORF. Black dots indicate individual data points. G. Overall survival for mice with D425 orthotopic xenografts in the murine cerebellum. sgControl mice (n=9) are shown in blue and sgASNSD1-uORF mice (n=10) are shown in red. P value is by a log-rank test. H. Brain MRIs at Day 22 post-injection for mice with sgControl orthotopic xenografts (#783, #788) or sgASNSD1-uORF orthotopic xenografts (#772, #775). Scale bars indicate relative scale.

This uORF encodes a conserved 96 amino acid sequence that spans four exons of the *ASNSD1* 5′-UTR and has recently been observed and annotated in prior non-canonical ORF discovery efforts (Figure 3C and Figures S3B-C).^41–43^ In humans, *ASNSD1* transcript expression is enriched in the cerebellum with preferential expression during early development, consistent with the location and onset of childhood medulloblastoma (Figures S3D-E).

To confirm its role in medulloblastoma cell viability, we performed CRISPR/Cas9 knockout validation experiments for ASNSD1-uORF across 5 MYC-driven and 4 non-MYC-driven medulloblastoma cell lines, as well as a larger set of 24 non-medulloblastoma cell lines. Loss of cell viability following knockout of ASNSD1-uORF was prominent in MYC-driven medulloblastoma cell lines, whereas 18/24 (75.0%) of non-MBL cell lines did not show a consistent phenotype (Figure 3E and **Table S3A**). Moreover, re-expression of the wild type ORF but not a start-site mutant rescued this phenotype (Figure 3F and Figure S3F), confirming the necessity of a protein-coding ASNSD1-uORF cDNA. In support of these observations, ectopic expression of ASNSD1-uORF led to a small but statistically significant increase in neural stem cell growth (9.8 vs. 7.9 doublings at 120 hours; p < 0.001, two-tailed Student’s T-test; Figures S3G-H).

We next investigated the role for ASNSD1-uORF in medulloblastoma *in vivo*. Consistent with its importance in medulloblastoma cell viability *in vitro*, knockout of ASNSD1-uORF prolonged overall survival for mice with orthotopic xenografts of D425 MYC-driven medulloblastoma cells (Figure 3G-H and Figure S3I). While editing efficiency was limited for *in vivo* knockouts, we observed that knockout allele fraction decreased, consistent with outgrowth of cells lacking allele knockout (Figure S3J). To probe a role in autochthonous medulloblastoma tumorigenesis, we performed *in utero* electroporation of ASNSD1-uORF cDNA in conjunction with cDNAs for cMYC and a dominant-negative p53 (DNp53) into the developing murine cerebellum. However, addition of ASNSD1-uORF to cMYC and DNp53 in this model did not alter mouse survival (Figure S3K-M).

### Elevated ASNSD1-uORF protein levels in medulloblastoma

Given the importance of ASNSD1-uORF in high-risk medulloblastoma, we next asked whether its abundance was increased in this disease. Indeed, ASNSD1-uORF displayed higher levels of RNA translation in MYC-driven cell lines by Ribo-seq (p = 0.0013, Figure 4A). Moreover, using targeted mass spectrometry with size selection, we observed a significant upregulation of ASNSD1-uORF protein level, but not other small proteins, in 10 MYC-driven compared to 5 non-MYC-driven medulloblastoma cell lines (p = 0.001, Figure 4B and Figure S4A). To validate these findings in patients, we leveraged publicly available mass spectrometry data for 45 pediatric medulloblastoma samples.^13^ In this historical dataset, we noted that ASNSD1-uORF appeared correlated with MYC in Group 3 tumors, though the analysis was underpowered (Figure S4B). Across all samples, high ASNSD1-uORF was also observed in samples in MYCN-high Group 4 tumors, where high MYC and high MYCN are mutually exclusive (Figure S4C-D). These results are consistent with the well-known overlap in MYC and MYCN function,^44^ as both may bind the same DNA motifs,^45^ dimerize with Max,^46^ and control similar downstream cellular programs.^47^ Therefore, we performed a merged analysis of ASNSD1-uORF protein levels in patient tumors with high levels of either MYC or MYCN, which revealed strong correlation between this uORF and the MYC family transcription factors (Pearson R = 0.47, p = 0.0009) (Figure 4C).^13^

**Figure 4:**
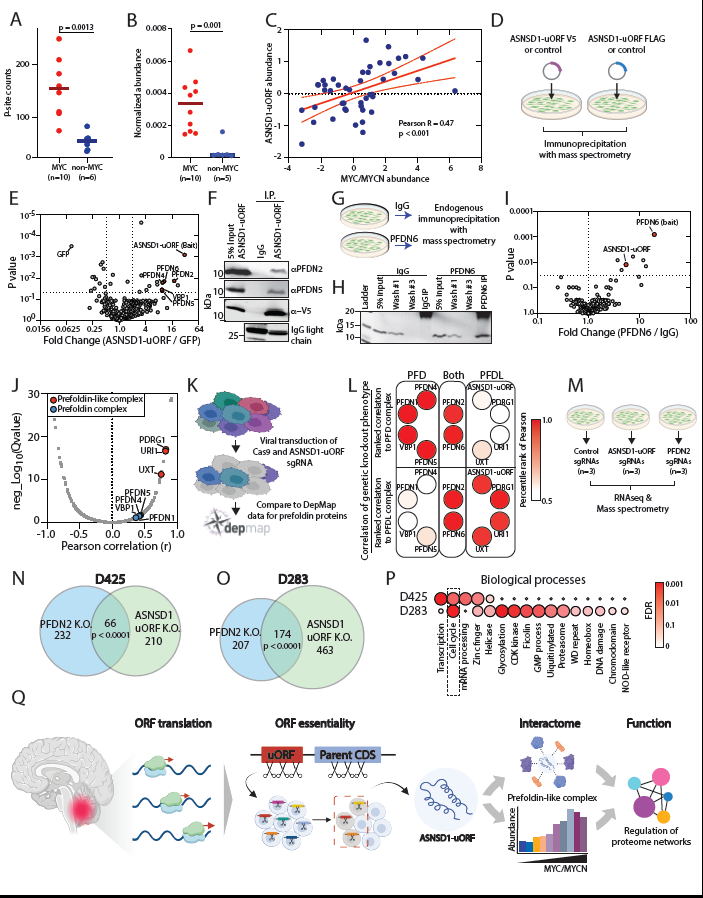
ASNSD1-uORF cooperates with the prefoldin-like complex in medulloblastoma. A. Abundance of ASNSD1-uORF translation across medulloblastoma cell lines using Ribo-seq data. Each dot reflects a cell line. P value by a two-tailed Student’s T-test. B. Protein abundance of ASNSD1-uORF in a cohort of MYC-driven (n=10) or non-MYC (n=5) medulloblastoma cell lines. P value by a two-tailed Student’s T-test. C. A scatter plot correlating ASNSD1-uORF protein abundance to protein abundance of MYC and MYCN in medulloblastoma patient samples (n=46) from the reanalyzed Archer *et al.* dataset.^13^ Correlation and p-value were determined by a Pearson R. D. A schematic showing the experimental design for ASNSD1-uORF co-immunoprecipitation from exogenous expression in HEK293T cells. E. A volcano plot showing enrichment of prefoldin and prefoldin-like complex proteins in ectopic ASNSD1-uORF co-immunoprecipitation in HEK293T cells. The X axis shows fold change of pull-down on a log2 scale. The Y axis shows the P value by a two-tailed Student’s t-test. F. A western blot showing validation of PFDN2 and PFDN5 pull-down with ASNSD1-uORF co-immunoprecipitation. G. A schematic showing the experimental design for endogenous co-immunoprecipitation with PFDN6. H. Western blot validation of PFDN6 pull-down in D425 cells. I. Mass spectrometry analysis of interacting partners with endogenous PFDN6 co-immunoprecipitation. The X axis shows fold change of pull-down on a log10 scale. The Y axis shows the P value by a two-tailed Student’s t-test. J. The correlation between ASNSD1-uORF protein abundance and prefoldin or prefoldin-like complex proteins from the reanalyzed Archer *et al.* medulloblastoma tissue samples (n=46).^13^ The X axis shows the Pearson correlation to ASNSD1 uORF. The Y axis shows the adjusted Q value. K. A schematic showing the experimental design for correlating ASNSD1-uORF knock-out phenotypes with knock-out phenotypes of prefoldin proteins. L. A heatmap showing the percentile rank of the Pearson correlation coefficient for loss of viability across 484 cancer cell lines following ASNSD1-uORF knockout or prefoldin/prefoldin-like gene knock-outs. M. A schematic showing the experimental design for RNA-seq and mass spectrometry experiments to functionally characterize ASNSD1-uORF N. Overlapping signatures of regulated proteins in mass spectrometry data for ASNSD1-uORF and PFDN2 knockout in D425. P value by a Fisher’s exact test. O. Overlapping signatures of regulated proteins in mass spectrometry data for ASNSD1-uORF and PFDN2 knockout in D283. P value by a Fisher’s exact test. P. Enriched biological processes identified in D425- or D283-signatures of proteins regulated by both PFDN2 and ASNSD1-uORF in mass spectrometry datasets. Q. A general model of non-canonical ORF translation in medulloblastoma.

We also measured ASNSD1-uORF protein levels across 23 non-medulloblastoma cell lines with matched CRISPR knockout data (as in Figure 3E, **Table S3A**) and observed that, while some cell lines lacking an essentiality phenotype expressed ASNSD1-uORF, medulloblastoma cell lines displayed both prominent protein expression and a loss-of-viability knockout phenotype (Figure S4E). Lastly, a reanalysis of mass spectrometry data for 504 solid tumor, non-medulloblastoma cancer cell lines^48^ demonstrated the greatest abundance of ASNSD1-uORF in MYCN-amplified neuroblastoma cell lines, consistent with MYC/MYCN regulation (Figure S4F). Taken together, our findings indicate that ASNSD1-uORF is a genetic dependency in high-risk medulloblastoma, which may be associated with its upregulation at the protein level in MYC or MYCN-driven pediatric cancers.

### ASNSD1-uORF functions coordinately with the prefoldin-like complex in medulloblastoma

To identify molecular mechanisms of ASNSD1-uORF in medulloblastoma, we pursued three strategies: protein-protein interactions, correlation of proteomic and genetic knockout signatures, and downstream molecular networks. First, we performed co-immunoprecipitation experiments for ectopically expressed ASNSD1-uORF followed by mass spectrometry (Figure 4D). Consistent with a prior report,^49^ we observed a striking enrichment for multiple members of the prefoldin complex, which we validated with western blots (Figure 4E-F). We further validated this interaction by using co-immunoprecipitation of endogenous prefoldin subunit 6 (PFDN6) in D425 cells (Figure 4G-H), which confirmed enrichment of endogenous ASNSD1-uORF protein (Figure 4I, Figure S4G, **Table S4A**).

Next, we sought to distinguish whether ASNSD1-uORF primarily operated in conjunction with the canonical prefoldin complex (PFD) or the more obscure prefoldin-like complex (PFDL) variant. The PFD is an evolutionarily conserved, hexameric protein chaperone complex thought to play an important role in the stability of nascent proteins.^26, 27^ Several clinicopathological studies have associated PFD components with cancer,^50–52^ including recent data that PFD proteins may be dysregulated in medulloblastoma.^16^ While the canonical PFD is embryonic lethal in mouse knockout models, the non-canonical PFDL – which retains only two of the six components of the PFD complex (PFDN2 and PFDN6) – may have only subtle murine knockout phenotypes (Figure S4H).

To place ASNSD1-uORF in the context of PFD or PFDL, we first used the Archer *et al.* medulloblastoma mass spectrometry dataset^13^ to correlate PFD or PFDL complex members to ASNSD1-uORF abundance. We observed that the PFDL-specific complex members are among the most highly correlated proteins with high statistical significance (p53 and DNA damage regulated 1 (PDRG1), URI1 prefoldin like chaperone (URI1) and ubiquitously expressed prefoldin like chaperone (UXT); Pearson correlations 0.756 - 0.826, Q values < 10^−12^ Figure 4J and **Table S4B**). By contrast, PFD complex-specific proteins were not significantly correlated with ASNSD1-uORF abundance. PDFL proteins were also significantly upregulated in MYC/MYCN driven medulloblastomas, similar to ASNSD1-uORF (Figure S4I). Next, we established that genetic knockout of PFDL proteins recapitulated the phenotype of ASNSD1-uORF knockout. Specifically, we used pooled cell culture to knockout ASNSD1-uORF in >400 barcoded PRISM cancer cell lines for dropout screening,^22^ and compared its pattern of genetic dependency to those of PFD and PFDL protein knockout in the same cell lines in the DepMap database (www.depmap.org) (Figure 4K, Figures S4J-K, and **Table S4C-F**). We found that members of the PFD and PFDL complexes readily clustered based upon the Pearson correlation of their knockout phenotype across the cell lines, and that ASNSD1-uORF was strongly associated with the PFDL but not the PFD complex (Figure 4L).

Knockout of ASNSD1-uORF or multiple prefoldin members did not impact the abundance of cytoskeletal proteins such as actin and tubulin, which have previously been suggested^53^ as downstream targets (Figure S4L). We therefore profiled transcriptomic and proteomic changes following knockout of ASNSD1-uORF or PFDN2 in D425 and D283 cells (Figure 4M and **Tables S4G-J**). Importantly, the protein abundance of the ASNSD1 parent CDS was not targeted by these gRNAs (Figure S4M). For both cell lines, we observed an overlapping proteomic signature of co-regulated proteins (Figure 4N-O), which demonstrated minimal change by RNA-seq (Figure S4N-O), confirming a post-transcriptional role for the prefoldin complex. Probing these sets of proteins further revealed consistent biological functional groups, with proteins related to cell cycle showing prominently (Figure 4P and **Table S4K**).

Collectively, these data support a role for ASNSD1-uORF within the PFDL complex in mediating cancer cell viability by coordinating downstream signatures of proteome regulation that may be relevant for medulloblastoma.

## Discussion

Here, we present a comprehensive analysis of the medulloblastoma translatome, generating matched Ribo-seq and RNA-seq data of 32 patient tissues and cell lines to enable the investigation of translated open reading frames in this disease. We show that medulloblastoma reproducibly translates over 6,700 non-canonical ORFs, which represent a previously unstudied layer of biology in this embryonal brain cancer (Figure 4Q). Using multiple CRISPR-Cas9 approaches to knockout over 2,000 ORFs, we broadly interrogate the contribution of non-canonical ORFs in cell survival across seven medulloblastoma cell lines. Overall, our results provide strong support for the growing community-wide interest in non-canonical ORFs as biological actors in both basic cell biology^20, 21, 30, 54–56^ and cancer pathophysiology.^22, 31, 57^ As such, our data argue for the inclusion of non-canonical ORFs in cancer genomics studies.

We particularly observe that a subset of uORFs function to maintain cancer cell survival. While early literature on uORFs has emphasized their importance only as regulators of mRNA translation,^18, 33, 58^ our efforts indicate that a sizable number of uORFs may operate as discrete biological actors. We are further able to pinpoint genetic dependency of 15 uORFs using high-density CRISPR tiling approaches, which provides high-resolution genetic evidence for uORF functionality in these cases. These data support the hypothesis that some uORFs are specific genetic dependencies in cancer even though the annotated, adjacent protein-coding CDS is not. Indeed, this hypothesis would suggest that some genes found to be dependencies by RNA interference screening – in which a full mRNA is downregulated – fail to score in CRISPR knockout data targeting the CDS. ASNSD1 points toward this: MYC-amplified medulloblastoma cell lines D458, D425 and D341 are among the most prominent hits in DEMETER shRNA data^59^ for ASNSD1, but do not score in the CRISPR-based DepMap (Figure S4P).

At the same time, we report the first example of molecular subtype-specific non-canonical ORF activity in childhood cancer. We focus on the role of the MYC family transcription factors, which we find may drive non-canonical ORF translation. Here, we establish a specific role for the ASNSD1-uORF as a medulloblastoma cancer dependency whose activity is linked to the MYC-family protein activity. Given the prominent role for MYC transcription factors in other cancer types, our observations that transcription factor amplification activates certain uORFs may have broader implications in cancer. To this end, we note that the example of ASNSD1-uORF is also more abundant in high-risk neuroblastoma cell lines, which may be due to impact on RNA translation by MYCN amplification.^60, 61^

Lastly, we describe a mechanism for ASNSD1-uORF within the poorly understood prefoldin-like complex, which is thought to play a role in protein homeostasis similar to that of the prefoldin complex, a related but distinct entity.^26, 53^ As such, our data reinforce a prior observation association ASNSD1-uORF with the prefoldin-like complex as well as emerging evidence that protein homeostasis via the prefoldin complex is dysregulated in medulloblastoma.^16^ While precise functions of the prefoldin-like complex remain incompletely understood, we observe that its impact on proteome regulation associates with specific, cancer-relevant biological functions, such as cell cycle. As a post-transcriptional mechanism of protein regulation, ASNSD1-uORF and the prefoldin-like complex lend additional evidence to observations that the medulloblastoma proteome deviates substantially from the transcriptome.^12, 13^

In summary, our findings exploit the known disease biology of medulloblastoma subtypes to provide cancer relevancy to the growing field of non-canonical ORFs and microproteins, providing context- and oncogene-specific consequences of non-canonical ORF translation. As such, our work provides additional rationale to investigate non-canonical ORFs and their translation as putative cancer target genes in medulloblastoma and other diverse malignancies.

## Acknowledgements

We thank Greg Newby and David Liu from the Broad Institute for their gracious insights with base editing experiments and providing the ABE8e-NRCH base editor. We thank Edmond Chan (Columbia University) for insights into chromatin immunoprecipitation experiments. We thank Ross Tomiano at the Taplin Mass Spectrometry Facility and Julian Mintseris at the Thermo-Fisher Center for Multiplexed Proteomics at the Harvard Medical School for assistance with mass spectrometry experiments. We thank Maura Berkeley and Zachary Herbert at the Dana-Farber Cancer Institute Molecular Biology Core Facility for assistance with next-generation sequencing. We thank the Dana-Farber Cancer Institute Center for Patient Derived Models for cell line support. We thank the Boston Children’s Hospital BioBank and the DFHCC Neurooncology Program Tissue and Data Bank for biobanking support. We thank the PRISM team at the Broad Institute for assistance with PRISM cell line screening and PRISM cell library sample preparation. We thank Joelle Straehla for sharing Med2112-mCherry-Luc and Med411-GFP-Luc cells. We thank Quang-De Nguyen, Amy Cameron and Murry Morrow at the Lurie Family Animal Imaging Center at the Dana-Farber Cancer Institute. J.R.P. acknowledges funding from the National Institutes of Health/National Cancer Institute (K08-CA263552-01A1), the Alex’s Lemonade Stand Foundation Young Investigator Award (#21-23983), the St. Baldrick’s Foundation Scholar Award (#931638), the DIPG/DMG Research Funding Alliance, the Cure ATRT Now Fund, the Musella Foundation for Brain Tumor Research, and a Collaborative Pediatric Cancer Research Awards Program/Kids Join the Fight award (#22FN23). T.R.G. acknowledges funding from the National Cancer Institute (1 R35 CA242457–01). K.L.L. and J.V. acknowledge support from the Pediatric Brain Tumor Foundation, the National Brain Tumor Society, and 3000 Miles to the Cure. S.v.H. acknowledges funding from Fonds Cancers (FOCA, Belgium), Stichting Reggeborgh (the Netherlands), and Bergh in het Zadel (the Netherlands). N.H. acknowledges funding from the European Union Horizon 2020 Research and Innovation Program (AdG788970), the Deutsche Forschungsgemeinschaft (SFB-1470 – B03), the Chan Zuckerberg Foundation (2019-202666), the Leducq Foundation (16CVD03), the British Heart Foundation and the Deutsches Zentrum für Herz-Kreislauf-Forschung (BHF/DZHK: SP/19/1/34461). P.B. acknowledges funding from the Isabel V. Marxuach Fund for Medulloblastoma Research, the Jared Branfman Sunflowers for Life Fund for Pediatric Brain and Spinal Cancer Research.

## Author contributions

Conceptualization: J.R.P., S.v.H., J.R.-O., and D.A.H.; methodology, J.R.P., D.A.H., S.v.H., J.R.-O, K.R.C., D.E.R., P.B.; validation, J.R.P., D.A.H., J.R.-O, R.M., I.Y.; formal analysis, J.R.P., D.A.H., J.R.-O, K.R.C.; investigation, J.R.P., D.A.H., J.R.-O., R.M., I.Y., K.N.Z., A.L.C., A.B., K.R.C., S.A.G.E., J.v.d.L., T.N.P., A.H., T.A., J.A.R., D.E.R., L.W., J.v.D.; resources, J.R.P., P.B., T.R.G., K.L.L., J.V., S.v.H., N.H.; data curation, J.R.P., D.A.H., J.R.-O; writing - original draft, D.A.H., J.R.-O., J.R.P., S.v.H.; writing - review & editing, D.A.H., J.R.-O., J.R.P., S.v.H. with input from all authors; visualization, J.R.P., J.R.-O., D.A.H.; supervision, J.R.P., S.v.H., P.B., T.R.G., N.H.; project administration: S.v.H. and J.R.P.; funding acquisition, J.R.P., T.R.G., N.H., K.L.L., P.B., and S.v.H.

## Inclusion and diversity

We support inclusive, diverse, and equitable conduct of research.

## Methods

### Data statement

Mouse xenografting experiments, our sample size of mice was predetermined based on the optimum number of animals needed to attain statistical significance of p<0.05 with a power level of 80 percent. For *in utero* electroporation, our sample size of 2-3 pregnant female mice to produce 12 electroporated murine pups per cohort reflects the known penetrance of tumor formation with cMYC and DNp53 with this technique,^62, 63^ and a sample size of 12 mice per cohort was designed to enable a statistical significance of p < 0.05 with a power level of 80 percent. Murine experiments were randomized and the investigators were blinded to allocation during experiments and outcome assessment.

### Cell lines and reagents

All parental cell lines were obtained directly from the American Type Culture Collection (ATCC, Manassas, VA), from the Bandopadhayay lab (MB002, D425, D458), Broad Institute Cancer Cell Line Encyclopedia (JIMT1, D384, R262, R256, UW228, HCC95, HCC15, SNU503, KYSE410, KYSE510, ONS76, RPE10-1), the Straehla lab (Med2112 and Med411) or from the Children’s Oncology Group (CHLA-259). H9-derived neural stem cells were obtained from Invitrogen (Invitrogen, cat# N7800-100). Cas9-derived cell lines were obtained from the Broad Institute. Cell lines were maintained according to established tissue culture media and conditions. HEK293T, D283Med (D283), D341, D384, D425, D458, DAOY, R262, R256, UW228, RPE10-1, JIMT1, MCF7, MDA-MB-231, HCC1806, HCC1954, HCC95, HCC15, A549, JURKAT, ES2, and MIAPACA2 cells were maintained in DMEM supplemented with 10% FBS (Invitrogen, Carlsbad, CA) and 1% penicillin-streptomycin (Invitrogen, Carlsbad, CA) in a 5% CO2 cell culture incubator. SNU503, HT29, KYSE410, KYSE510, ONS76, A375, HS294T, and LOXIMVI cells were maintained in RPMI 1640 (Invitrogen, Carlsbad, CA) supplemented with 10% FBS and 1% penicillin-streptomycin in a 5% CO2 cell culture incubator. CHLA-259 cells were maintained in IMDM (Invitrogen, Carlsbad, CA) supplemented with 20% FBS and 1% penicillin-streptomycin in a 5% CO2 cell culture incubator. CHLA-02-ATRT, CHLA-05-ATRT, CHLA-06-ATRT, CHLA-01-MED, CHLA-01-MEDR, H9-derived NSCs, Med2112 (expressing mCherry and luciferase), Med411 (expressing GFP and luciferase) and MB002 cells were maintained in Tumor Stem Media comprised of DMEM/F12 (1:1) with Neurobasal-A medium (Invitrogen, Carlsbad, CA) and supplemented with HEPES (1M, 0.1% final concentration; Invitrogen, Carlsbad, CA), sodium pyruvate (1mM final concentration; Invitrogen, Carlsbad, CA), MEM non-essential amino acids (0.1mM final concentration; Invitrogen, Carlsbad, CA), GlutaMax (1x final concentration; Invitrogen, Carlsbad, CA), B27 supplement minus vitamin A (1x final concentration; Invitrogen, Carlsbad, CA), human EGF (20ng/mL; Shenandoah Biotech), human FGF-basic-154 (20ng/mL; Shenandoah Biotech), and Heparin solution 0.2% (2ug/mL final concentration, StemCell Technologies). H9-derived NSC cells were cultured on GelTrex-coated tissue culture plates (ThermoFisher). Cell lines were routinely verified via STR genotyping and tested for mycoplasma contamination using the Lonza MycoAlert assay (Lonza). Below is a list of cell line details:

| Cell line | Source | Catalog number | Culture media | RRID |
| --- | --- | --- | --- | --- |
| CHLA-259 | Children's Oncology Group | CHLA-259 | IMDM, 20% FBS, 1% PS | CVCL_M148 |
| MB002 | Bandopadhayay lab |  | Tumor Stem Media | CVCL_VU79 |
| H9-NSCs | Invitrogen | N7800-100 | Tumor Stem Media | CVCL_IU37 |
| HEK293T | ATCC | CRL-3216 | DMEM, 10% FBS, 1% PS | CVCL_0063 |
| D283Med | ATCC | HTB-185 | DMEM, 10% FBS, 1% PS | CVCL_1155 |
| D341 | ATCC | HTB-187 | DMEM, 10% FBS, 1% PS | CVCL_0018 |
| JIMT1 | CCLE |  | DMEM, 10% FBS, 1% PS | CVCL_2077 |
| D425 | Bandopadhayay lab |  | DMEM, 10% FBS, 1% PS | CVCL_1275 |
| D458 | Bandopadhayay lab |  | DMEM, 10% FBS, 1% PS | CVCL_1161 |
| D384 | CCLE |  | DMEM, 10% FBS, 1% PS | CVCL_1157 |
| DAOY | ATCC | HTB-186 | DMEM, 10% FBS, 1% PS | CVCL_1167 |
| R262 | CCLE |  | DMEM, 10% FBS, 1% PS | CVCL_VU83 |
| R256 | CCLE |  | DMEM, 10% FBS, 1% PS | CVCL_DG09 |
| UW228 | CCLE |  | DMEM, 10% FBS, 1% PS | CVCL_8585 |
| RPE10 | CCLE |  | DMEM, 10% FBS, 1% PS | CVCL_4388 |
| MCF7 | ATCC | HTB-22 | DMEM, 10% FBS, 1% PS | CVCL_0031 |
| MDA-MB-231 | ATCC | CRM-HTB-26 | DMEM, 10% FBS, 1% PS | CVCL_0062 |
| HCC1806 | ATCC | CRL-2335 | DMEM, 10% FBS, 1% PS | CVCL_1258 |
| HCC1954 | ATCC | CRL-2338 | DMEM, 10% FBS, 1% PS | CVCL_1259 |
| HCC95 | CCLE |  | DMEM, 10% FBS, 1% PS | CVCL_5137 |
| HCC15 | CCLE |  | DMEM, 10% FBS, 1% PS | CVCL_2057 |
| A549 | ATCC | CRM-CCL-185 | DMEM, 10% FBS, 1% PS | CVCL_0023 |
| Jurkat | ATCC | TIB-152 | DMEM, 10% FBS, 1% PS | CVCL_0065 |
| ES2 | ATCC | CRL-1978 | DMEM, 10% FBS, 1% PS | CVCL_AX39 |
| MIAPACA2 | ATCC | CRM-CRL-1420 | DMEM, 10% FBS, 1% PS | CVCL_0428 |
| SNU503 | CCLE |  | RPMI1640, 10% FBS, 1% PS | CVCL_5071 |
| HT29 | ATCC | HTB-38 | RPMI1640, 10% FBS, 1% PS | CVCL_0320 |
| KYSE410 | CCLE |  | RPMI1640, 10% FBS, 1% PS | CVCL_1352 |
| KYSE510 | CCLE |  | RPMI1640, 10% FBS, 1% PS | CVCL_1354 |
| ONS76 | CCLE |  | RPMI1640, 10% FBS, 1% PS | CVCL_1624 |
| A375 | ATCC | CRL-1619 | RPMI1640, 10% FBS, 1% PS | CVCL_0132 |
| HS294T | ATCC | HTB-140 | RPMI1640, 10% FBS, 1% PS | CVCL_0331 |
| LOXIMVI | Millipore Sigma | SCC201 | RPMI1640, 10% FBS, 1% PS | CVCL_1381 |
| CHLA-02-ATRT | ATCC | CRL-3020 | Tumor Stem Media | CVCL_B045 |
| CHLA-05-ATRT | ATCC | CRL-3037 | Tumor Stem Media | CVCL_AQ41 |
| CHLA-06-ATRT | ATCC | CRL-3038 | Tumor Stem Media | CVCL_AQ42 |
| CHLA-01-MED | ATCC | CRL-3021 | Tumor Stem Media | CVCL_B044 |
| CHLA-01-MEDR | ATCC | CRL-3034 | Tumor Stem Media | CVCL_N534 |
| Med2112-mCherry-Luc | Straehla lab |  | Tumor Stem Media | NA |
| Med411-GFP-Luc | Straehla lab |  | Tumor Stem Media | NA |

### Tissue samples

21 human medulloblastoma tissue samples were obtained from the Boston Children’s Hospital BioBank and the Dana-Farber Harvard Cancer Center Neuro-oncology Program and Tumor BioBank. Patient samples were acquired with the informed consent of DFCI protocol 10-417. Four human medulloblastoma tissue samples were obtained from the Princess Máxima Center biobank under approval from the Medical Ethics Committee of the Erasmus Medical Center (ID number, MEC-2016-739). All samples were de-identified prior to use for research.

### Immunoblot Analysis

Cells were grown to 70-80% confluence, collected by scraping the tissue culture dish and washed once in 1x PBS. They were then lysed in RIPA lysis buffer (Sigma-Aldrich, St. Louis, MO) with 1x HALT protease inhibitor (Thermo Fisher Scientific, Waltham, MA) and homogenized by chilling them on ice for 15 minutes. Cellular proteins were separated by centrifugation for 15 minutes at 13,200 RPM and supernatant was saved. Protein lysate yields were determined using bicinchoninic acid (BCA), and appropriate volumes of lysate were prepared for immunoblotting by boiling in a 1x sample loading buffer at 95C for 5 minutes. Tris-Glycine 10-20% or Bis-Tris 4-12% SDS-PAGE gels were run at 4℃ and proteins were transferred onto nitrocellulose membranes using 15 Volts for 7 minutes via the iBlot-2 system (Thermo Fisher Scientific, Waltham, MA). The membrane was then blocked for 1 hour in LICOR Odyssey blocking buffer and incubated at 4℃ with the appropriate antibody overnight. The blot was then washed 4 times with 1x TBS with 0.1% Tween20 and incubated with fluorophore-specific IRDye secondary antibodies (LI-COR, Lincoln, NE) and imaged on a LI-COR Odyssey machine.

### Immunoblot antibodies used

Immunoblot antibodies used:
| Antibody | Type | Species | Monoclonal/<br>Polyclonal | Dilution | Indication | Catalog<br>Number | Vendor | Conditions | RRID |
| --- | --- | --- | --- | --- | --- | --- | --- | --- | --- |
| V5<br>(D3H8Q) | Primary | Rabbit | Monoclonal | 1:2500 | Western blot,<br>co-IP | 13202S | Cell Signaling<br>Technology | 4C overnight | AB_26874<br>61 |
| V5 [SV5-Pk1] | Primary | Mouse | Monoclonal | 1:2500 | Western blot | ab27671 | Abcam | 4C overnight | AB_47109<br>3 |
| GFP | Primary | Rabbit | Polyclonal | 1:2500 | Western blot | 2555S | Cell Signaling<br>Technology | 4C overnight | AB_10692<br>764 |
| PFDN1 | Primary | Rabbit | Polyclonal | 1:1000 | Western blot | HPA00649<br>9 | Millipore<br>Sigma | 4C overnight | AB_10795<br>96 |
| PFDN2 | Primary | Rabbit | Polyclonal | 1:500 | Western blot | HPA02870<br>0 | Millipore<br>Sigma | 4C overnight | AB_10603<br>983 |
| PFDN5 | Primary | Rabbit | Polyclonal | 1:500 | Western blot | HPA00858<br>7 | Millipore<br>Sigma | 4C overnight | AB_10795<br>97 |
| PFDN6 | Primary | Rabbit | Polyclonal | 1:1000 | Western blot,<br>co-IP | HPA04303<br>2 | Millipore<br>Sigma | 4C overnight | AB_26782<br>78 |
| Alpha-<br>tubulin | Primary | Mouse | Monoclonal | 1:2000 | Western blot | ab7291 | Abcam | 4C overnight | AB_22411<br>26 |
| Beta-tubulin | Primary | Rabbit | Polyclonal | 1:2000 | Western blot | 2146S | Cell Signaling Technology | 4C overnight | AB_2210545 |
| Gamma-tubulin | Primary | Rabbit | Polyclonal | 1:1000 | Western blot | A302-631A-M | Bethyl Laboratories | 4C overnight | AB_2780661 |
| HSP90 | Primary | Rabbit | Monoclonal | 1:1000 | Western blot | 4877S | Cell Signaling | 4C overnight | AB_2233307 |
| Vinculin | Primary | Rabbit | Monoclonal | 1:1000 | Western blot | ab219649 | Abcam | 4C overnight | AB_2819348 |
| GAPDH | Primary | Rabbit | Monoclonal | 1:2000 | Western blot | 2118L | Cell Signaling Technology | 4C overnight | AB_561053 |
| Beta Galactosidase | Primary | Rabbit | Polyclonal | 1:2000 | Western blot | Ab616 | Abcam | 4C overnight | AB_305327 |
| Beta-Actin | Primary | Mouse | Monoclonal | 1:4000 | Western blot | A5316 | Sigma-Aldrich | 4C overnight | AB_476743 |
| Goat anti-mouse secondary | Secondary | Goat | N/A | 1:5000 | Western blot | 926-32210 | LI-COR | 20C for 1 hour | AB_621842 |
| Goat anti-rabbit secondary | Secondary | Goat | N/A | 1:5000 | Western blot | 926-68021 | LI-COR | 20C for 1 hour | AB_10706309 |

### RNA isolation and cDNA synthesis

Total RNA was isolated using Qiazol and an miRNeasy Kit (Qiagen, Hilden, Germany) with DNase I digestion according to the manufacturer’s instructions. RNA integrity was verified on an Agilent Bioanalyzer 2100 (Agilent Technologies, Palo Alto, CA). cDNA was synthesized from total RNA using Superscript III (Invitrogen, Carlsbad, CA) and random primers (Invitrogen, Carlsbad, CA).

### Ribosome profiling

Ribo-seq for human tissue samples was performed according to the protocol described in Palomar-Siles et al.^64^ Ribo-seq for cancer cell lines was performed based upon the protocol by McGlincy et al.^65^ with modifications as described below. Briefly, cells were grown to 60-70% confluence prior to collection. After collection, all cell pellets were washed once in 1x PBS, re-pelleted by centrifugation, and lysed in lysis buffer (20mM Tris HCl, 150mM NaCl, 5mM MgCl2, 1mM dithiotrietol, 0.05% NP-40, 25U/mL Turbo-DNase I (Invitrogen), 2ug/mL cycloheximide). After clearing the lysate and recovering the supernatant, RNA abundance was determined by measuring the A260. 2.5U/ug of RNase I was added to an appropriate volume of lysate and incubated at 22C for 45 minutes without shaking. The RNase I was then quenched with 1U/uL of Superase RNase Inhibitor (Ambion). RNA from ribosome protected fragments were recovered using a 1M sucrose cushion with ultracentrifugation (55,000 RPM, 4C, 2 hours), and rRNA was depleted using the siTOOLS human RiboPool kit according to manufacturer’s instructions (siTOOLS Biotech, Germany). Ribosome protected fragments were then denatured using a 1:1 mixture with 2x sample loading buffer (98% v/v formamide, 10mM EDTA, 300ug/mL bromophenol blue) at 95C for 3 minutes, and further purified using size selection from a 15% TBE-Urea gel (200V for 65 minutes). The 26 – 32 nucleotide band was cut from the gel, RNA extracted by freezing gel slices in 400uL RNA gel extraction buffer (300mM NaOAc, 1mM EDTA, 0,25% v/v SDS), and rotating at room temperature for 5-6 hours. RNA was precipitated with 500uL isopropanol and 2.0uL GlycoBlue at −20C overnight; pellets were washed once in chilled 70% ethanol, and subjected to end-repair with T4 PNK (Lucigen, 37C for 1 hr). End-repaired RNA was cleaned up with the RNA Clean and Concentrator kit (Zymo), ligated to a 3′ linker (sequence below, 6.67% w/v PEG-8000, 6.67 mM dithiotrietol, 1x T4 RNL2 Truncation buffer, 6.67 U/uL R4 RNA ligase 2 Deletion mutant, 0.33 U/uL T4 RNA ligase I) for 3 hours at room temperature. Linker reactions were removed with 5′ deadenylase (New England Biolabs) and Rec J Exonuclease (NEB), and cDNA was generated with EpiScript RT enzyme (Lucigen, 50C for 30 minutes) followed by reaction clean up with exonuclease I (Lucigen, 37C for 30 minutes), RNase I/Hybridase (Lucigen, 55C for 5 minutes) and the Oligo Clean and Concentrator Kit (Zymo). cDNA was mixed 1:1 with 2x sample loading buffer, boiled, and purified with a 10% TBE-Urea gel (70 minutes, 175V). The product between 70 – 90 nucleotides was excised from the gel, and DNA was extracted with 450uL DNA extraction buffer (300mM NaCl, 10mM Tris, 1mM EDTA, 0.02% SDS) with a flash-freeze on dry ice (30 minutes) and rotation at 22C for 6 hours. DNA was precipitated with 700uL isopropanol and 2uL GlycoBlue at −80C overnight followed by centrifugation at 14,500 RPM for 45 minutes at 4C. DNA pellets were washed once in 80% ethanol and pellets were air-dried and dissolved in 11uL of water, which was then circularized with the addition of 9uL of CircLigase I mix (1M betaine, 1x CircLigase Buffer (Lucigen), 2.5mM MnCl2, 50uM ATP, 5U/uL CircLigase I (Lucigen)) at 60C for 3 hours with heat inactivation at 80C for 10 minutes. Circularized cDNA was quantified using quantitative real-time PCR (10uL of 2x SYBR-Green mastermix (Thermo), 2uL of cDNA, 6uL water, 1uL forward and reverse primer each) for twenty cycles, using the following PCR primers (JRP_qPCR-ribo-F2 primer: CAGAGTTCTACAGTCCGACGAT; JRP_qPCR-ribo-R2 primer: AGACGTGTGCTCTTCCGATCT). Library PCR amplification was performed with 10uL of 2x Phusion HiFi master mix (New England Biolabs), 8uL of cDNA sample, 1uL of the forward library primer (AATGATACGGCGACCACCGAGATCTACACGTTCAGAGTTCTACAGTCCGACG) and 1uL of the appropriate barcorded reverse primer (**Table S4L**). PCR reactions were run with the following cycle conditions: 98C for 1 minute, followed by 12-15 cycles of 94C for 16 seconds, 55C for 6 seconds, and 65C for 11 seconds, with a final extension of 65C for 1 minute. PCR products were mixed with 6x gel loading buffer and size-selected on a 8% TBE gel, 100V for 75 minutes. The product at ∼150 bps was gel-excised, placed in 400uL of DNA extraction buffer, flash frozen on dry ice for 30 minutes, thawed at 22C for 6 hours on a rotating platform, and DNA was precipitated with 700uL of isopropanol with 2uL of GlycoBlue overnight at −80C. Samples were then centrifuged at 14,500 RPM for 45 minutes; DNA pellets were washed once in 80% ethanol, air-dried, and dissolved in 18uL of 5mM Tris. Samples were quantified by DNA Qubit (Thermofisher) and library size was confirmed using an Agilent Bioanalyzer HS DNA High Sensitivity Kit (Agilent). Libraries were sequenced at the Dana-Farber Molecular Biology Core Facility on an Illumina NovaSeq 6000.

### RNA sequencing

Matched RNA sequencing for all samples was performed by removing 1/3^rd^ of the sample lysate from the ribosome profiling sample and placing it in 400uL Trizol. RNA was then extracted using the Qiagen RNAeasy kit (Qiagen) according to the manufacturer’s instructions. RNA abundance was quantified using spectrophotometry via Nanodrop as well as RNA Qubit (Thermofisher). RNA samples were submitted to the Dana-Farber Molecular Biology Core Facility for mRNA sequencing using the Roche Kapa mRNA Hyper Prep kit (Roche, Basel, Switzerland) with samples sequenced on an Illumina NextSeq or NovaSeq. RNA samples from the Princess Maxima Center were processed through the Princess Maxima Center Diagnostics core facility according to institutional protocols.

### Analysis of RNA-seq data for sample clustering and gene set enrichment analysis

The raw RNA-seq reads from cell lines and tissue samples were subjected to quality control and read trimming using TrimGalore v0.6.6^66^, which internally employs Cutadapt v3.4 ^67^ for adapter removal and FastQC v0.11.9 (https://www.bioinformatics.babraham.ac.uk/projects/fastqc/) for quality assessment. using standard parameters for paired-end reads.

Trimmed and filtered reads were aligned to human reference genome hg38 using STAR v2.7.8a ^68^ in the two-pass mapping mode, with genome annotation provided in GTF format (Ensembl release 102). Default STAR settings were used, with the following modified parameters: *--outFilterType BySJout --outSAMunmapped Within --outSAMattributes NH HI AS nM NM MD jM jI MC ch --outSAMstrandField intronMotif --outSAMtype BAM Unsorted --outFilterMismatchNmax 6 --alignSJoverhangMin 10 --outFilterMultimapNmax 10 --outFilterScoreMinOverLread 0.75*.

Counts for annotated CDS regions were obtained using featureCounts v2.0.2^69^ with genome annotation provided in GTF format (Ensembl release 102), and CDS regions used as the counting feature in paired-end mode. To improve read counting for junctions, the *–J* option was used with reference sequences for transcripts provided in FASTA format (GRCh38, Ensembl release 102).

CDS read counts from the cell line samples (annotated as either MYC high or MYC low) were used as input for DESeq2^70^ to perform principal component analysis and differential expression analysis, using the default DESeq2 workflow and MYC status (MYC high vs MYC low) as contrasting variable.

Gene ontology (GO), hallmark, KEGG, and Reactome gene sets were obtained from the MSigDB ^71, 72^ database using the msigdb R package^73^, and were used as query gene sets. A list of log2 fold change values, obtained from the DESeq2 output, was used as input for gene set enrichment analysis using the fgsea R package^74^. Gene set enrichment analysis was performed separately for each of the gene set categories (GO:CC, GO:BP, GO:MF, Hallmark, Reactome, KEGG). Gene sets with an adjusted P-value < 0.05 and a normalized enrichment score > 0 were considered significantly over-enriched in MYC-driven compared to non-MYC-driven samples.

### Processing of RNA-seq data for gene-level translational efficiency calculation

To facilitate comparison with ribo-seq data and calculate translational efficiency values, the RNA-seq reads were reprocessed using different alignment and filtering parameters as described below.

The raw RNA-seq reads were subjected to quality control and read trimming using TrimGalore. Only the first reads of the read pairs were used, to imitate single-end ribosome profiling reads. The RNA-seq reads were hard-trimmed to 29-mers using Cutadapt with the *--hardtrim5* option. Then, TrimGalore was run on the trimmed reads with options set to remove Ns (*--trim-n*) and retain reads with a minimum length of 25 bp (*--length 25*). FastQC was executed within TrimGalore to remove low quality reads.

To eliminate reads corresponding to contaminants such as tRNA, rRNA, snRNA, snoRNA, and mtDNA, Bowtie2 (v2.4.2)^75^ was executed with standard parameters and option *--seedlen=25* to align the reads to a custom reference database containing sequences of these contaminants. The unaligned reads, i.e., those not mapping to any of the contaminants, were output to a gzipped FASTQ file for further processing.

The filtered reads were aligned to reference genome GRCh38 using STAR v2.7.8a with options *--outFilterMismatchNmax 2 --outFilterMultimapNmax 20 --outSAMattributes All --outSAMtype BAM SortedByCoordinate --quantMode GeneCounts --limitOutSJcollapsed 10000000 --outFilterType BySJout --alignSJoverhangMin 1000*, using the MANE Select v1.0^76^ transcript annotation, supplied in a GTF file, as reference annotation.

To quantify reads aligning to annotated CDS features, featureCounts was used with the options *--J, --t “CDS”, --g “gene_id”,* resulting in CDS counts summarized on gene-level. Annotations and sequences for reference transcripts for GRCh38 / Ensembl release 102 were provided in FASTA and GTF files, respectively.

### Ribosome profiling read alignment and processing

Raw ribosome profiling reads were trimmed and filtered using TrimGalore with the following options: *--gzip --length 25 --trim-n*. Contaminant reads were filtered out with Bowtie2 with the option *--seedlen=25*, using a custom index containing tRNA, rRNA, snRNA, snoRNA, and mtDNA sequences. Filtered ribo-seq reads were aligned to reference genome GRCh38 using STAR v2.7.8a with options *--outFilterMismatchNmax 2 --outFilterMultimapNmax 20 --outSAMattributes All --outSAMtype BAM SortedByCoordinate --quantMode GeneCounts --limitOutSJcollapsed 10000000 --outFilterType BySJout --alignSJoverhangMin 1000*, using GRCh38 / Ensembl release 102 reference annotation provided in GTF file. Annotated CDS features were quantified using featureCounts with the options *--J --t “CDS” --g “gene_id”,* with Ensembl release 102 annotation provided in GTF format and GRCh38 / Ensembl release 102 transcript sequences provided in FASTA format. We then used RiboseQC^77^ provided with Ensembl release 102 transcript annotation in GTF format to assess data quality and quantify P-site positions in the aligned ribo-seq reads in all samples.

### Calculating translational efficiency values

Translational efficiency values for annotated genes were calculated using gene-summarized RNA-seq and Ribo-seq CDS read counts in cell line samples. To ensure that the genes used for TE calculation showed robust expression in both ribo-seq and RNA-seq data, genes with fewer than 128 read counts on average across all samples in either RNA-seq or ribo-seq were removed. To make the RNA-seq and ribo-seq read counts comparable, they were first converted to TPM values. The TE for each gene was then calculated as the ratio of TPM(ribo-seq) over TPM(RNA-seq). Non-real values resulting from divisions by zero were set to 0. To plot the densities of the translational efficiency values for all genes in MYC-driven and non-MYC samples, the TE values were log2-transformed and centered by subtracting the TE value of each gene in each sample by the median TE of that gene across all samples.

### P-site quantification and determining translated ORFs

To quantify ribo-seq P-sites on an ORF level, we generated BED files that contain all possible P-site positions for annotated as well as non-canonical ORFs. We used a GTF file containing MANE Select transcript definitions^76^ (matching the Ensembl annotations in version hg38) to obtain annotations for annotated CDS regions, and a custom GTF file containing merged definitions from GENCODE Phase 1 ORFs^17^ and our prior custom cancer ORFeome^17, 22^ for non-canonical ORFs. A custom Python script was used to generate ‘reference’ BED files containing the coordinates of all potential P-site positions for each ORF, annotated by frame (p0, p1, or p2) in each codon. Incomplete proteins were excluded using provided annotation files (see Data Availability statement).

P-site coordinates and counts in each sample were extracted from RiboseQC output files and stored in BED files. Bedtools intersect v2.25.0^78^ was used to overlap detected P-sites with the ‘reference’ P-sites using the options ‘-wa -wb -header -f 1.00 -s’. For each sample, the resulting BED files contained the P-site coordinates, counts, and ORF names (annotated and non-canonical) of overlapping ‘reference’ P-sites.

The resulting intersected BED were then used to generate a matrix of P-site counts per ORF in each sample. To construct this matrix, we first calculated the frame with the highest P-site fraction for each ORF in a given sample. We then added the total P-site count of the dominant frame of each ORF to the P-site count matrix.

To identify translated ORFs, P-site counts were converted to TPM-like count values (P-sites per million, or PPM). First, P-sites for each ORF were divided by the ORF length in kb to calculate P-sites per kb (PPK). Per-million scaling factors for each sample were calculated by dividing the sum of each sample’s ppk values by 1,000,000. Each ORF’s PPM value was then calculated by dividing the ORF’s PPK by the sample’s scaling factor. To define a PPM cutoff for determining translation, the density of log2-transformed PPM values was plotted and visually inspected. There was a clear bimodal distribution, so we selected a cutoff value between the low and high distributions, which corresponded to a PPM value of 1. Translated ORFs were then defined as ORFs with a PPM > 1 in at least 5 samples.

### Identifying differentially translated ORFs

The matrix with raw ORF P-site counts for the cell line samples was loaded into R and used as input for DESeq2 to perform principal component analysis and differential expression analysis, using the default DESeq2 workflow, and using MYC status (MYC-driven vs non-MYC) as contrasting variable. The volcano plot showing differentially translated ORFs between MYC-driven and non-MYC samples was generated using the EnhancedVolcano R package.^79^ ORFs were sorted by p-value, and top 5 upregulated (log2 fold change > 2) and top 5 downregulated (log2 fold change < −2) were highlighted.

### ORF-level translational efficiency analysis

To obtain ORF-level RNA-seq read counts, we used Salmon v1.8.0,^80^ with Bowtie2-filtered raw RNA-seq reads as input (see section <u>Processing of RNA-seq data for gene-</u> <u>level translational efficiency calculation)</u>. A custom Salmon index was generated based on a custom GTF file containing the merged set of annotated MANE transcripts as well as non-canonical GENCODE Phase1^17^ (https://www.gencodegenes.org/pages/riboseq_orfs/) and ORFeome definitions (**Table S1L**). Briefly, CDS regions were extracted from the custom GTF file and stored in a separate, cleaned up GTF file with transcript IDs set to match ORF IDs, since Salmon uses transcript IDs to differentiate between features. The CDS GTF file was cleaned up with We ran Salmon with the following parameters: *‘salmon quant --libtype “A” --validateMappings --gcBias --numGibbsSamples 30’*.

We loaded the matrices with ORF-level RNA-seq counts and P-site counts for the cell line samples into R, and removed ORFs with fewer than 128 counts on average across all samples in either RNA-seq or ribo-seq. We calculated TPM and PPM values for the remaining ORFs. Translational efficiency for each ORF was calculated as the ratio of TPM(Ribo-seq) over TPM(RNA-seq). Non-real values resulting from divisions by zero were set to 0. TE values were log2-transformed and scaled to perform principal component analysis. The full code can be found at: https://github.com/damhof/hofman_et_al_2023_seq

### Determination of infection conditions for CRISPR pooled screens

Optimal infection conditions were determined in each cell line in order to achieve 30-50% infection efficiency, corresponding to a multiplicity of infection (MOI) of ∼0.5 −1. Spin-infections were performed in 12-well plate format with 3 x 10e6 cells each well. Optimal conditions were determined by infecting cells with different virus volumes with a final concentration of 4 ug/mL polybrene. Cells were spun for 2 hours at 1000 g at 30 degrees. Approximately 24 hours after infection, cells were trypsinized and approximately 2×10e5 of R262, UW228, ONS76, D458, D425, D283, or D341 cells from each infection were seeded in 2 wells of a 6-well plate, each with complete medium, one supplemented with 1.5ug/mL of puromycin. Cells were counted 4-5 days post selection to determine the infection efficiency, comparing survival with and without puromycin selection. Volumes of virus that yielded ∼30 −50% infection efficiency were used for screening.

### Primary and validation CRISPR Pooled Proliferation Screens

The lentiviral barcoded library used in the primary screen contains 26,819 sgRNAs and the validation library contains 6,557 gRNAs targeting selected regions of the ORFs, which were designed using the CRISPick program (https://portals.broadinstitute.org/gppx/crispick/public) from Broad Institute Genomic Perturbation Platform, using settings for the reference genome Human GRCh38 (Ensembl v.108) for “CRISPRko” with enzyme “SpyoCas9 (NGG)” with the following modifications:

- Each ORF and parental CDS were targeted by up to 8 gRNAs where possible. A distribution of the number of gRNAs per target is displayed in **Table S2A**.
- For ORFs with >= 2 exons, the best gRNA design was selected for each exon to a maximum of 8 gRNAs. For ORFs with >2 but <8 exons, the remaining gRNAs were selected as the top picks from any exon.
- The spacing requirement for gRNA separation was reduced to 1% across the total target length for ORFs and maintained at 5% for parental CDSs.
- A 2:1 on-target to off-target ratio was employed.
- For the validation library, ORFs were targeted with a maximum of 24 gRNAs per exon, 5′UTR and 3′UTRs with a maximum of 12 gRNAs per UTR region, up to 3 introns with 6 gRNAs per intron, the upstream genome promoter region with 6 gRNAs (defined as within 1000 basepairs of the transcript start site), and up to 3 parental CDS exons with 8 gRNAs per exon.
- Both libraries employed a common set of 503 non-targeting gRNAs without genome cutting, and 497 non-targeting gRNAs with genome cutting for negative controls. The primary library had 1694 positive control pan-lethal gRNAs. The validation library had 527 positive control pan-lethal gRNAs.

Genome-scale infections were performed in three replicates with the pre-determined volume of virus in the same 12-well format as the viral titration described above, and pooled 24 h post-centrifugation. Infections were performed with enough cells per replicate, in order to achieve a representation of at least 500 cells per gRNA (for primary screen) or 1000 cells per gRNA (for validation screen) following puromycin selection (∼1.5×10e7 surviving cells). Approximately 24 hours after infection, all wells within a replicate were pooled and were split into T225 flasks. 24 hours after infection, cells were selected with puromycin for 7 days to remove uninfected cells. After selection was complete, 1.5-2×10e7 of cells were harvested for assessing the initial abundance of the library. Cells were passaged every 3-4 days and harvested ∼14 days after infection. For all genome-wide screens, genomic DNA (gDNA) was isolated using Midi or Maxi kits for the validation screens gDNA was isolated using Midi kits according to the manufacturer’s protocol (Qiagen). PCR and sequencing were performed as previously described.^81, 82^ Samples were sequenced on a HiSeq2000 or NextSeq (Illumina). For analysis, the read counts were normalized to reads per million and then log2 transformed. The log2 fold-change of each sgRNA was determined relative to the initial time point for each biological replicate.

### Analysis of CRISPR screening data

CRISPR data was transformed into log2 fold change values computed between the day 14 timepoint and the input plasmid DNA. All values were then normalized to the positive control gRNAs in the following way: for each cell line, the gRNAs targeting parental_poscon genes were averaged. This geometric mean of the poscons was scaled to equal −1. This was accomplished by dividing individual gRNA values by the poscon mean, multiplied by −1 to retain a negative value to represent gRNA drop-out. The equation is as follows: (gRNA/average_poscon)*-1. A “hit” was defined as a non-canonical ORF that had at least 2 gRNAs with a normalized abundance of less than or equal to −1.0 at the day 14 timepoint in the primary screen. For uORFs, uoORFs, and dORFs, the comparison between the non-canonical ORF and the parental CDS should demonstrate a differential effect (delta_ORF-CDS effect) of less than or equal to −0.3 to yield a potential differential dependency. uORFs, uoORFs and dORFs were further assessed by comparing the absolute number of gRNAs with a normalized abundance of less than or equal to −1.0 to the absolute number of parental CDS gRNAs with a normalized abundance of less than or equal to −1.0.

### Assessment of toxicity of Cas9 activity at gene promoters

To assess Cas9 toxicity when targeting uORFs located near to the gene promoter, the primary screen further targeted 120 pan-lethal positive control genes known to have a uORF as well as 82 pan-lethal positive control genes with no known uORF. For the latter, a 150 bp segment of the gene 5′UTR was targeted with gRNAs. The data were analyzed as described above to estimate the potential impact of Cas9 genome toxicity at the promoters of genes. Figure S2K provides additional details.

### Analysis of CRISPR validation screen

The validation screen targeted 44 uORFs, 6 uoORFs, 10 lncRNA-ORFs, and their associated parental CDS and genomic regions (**Table S2I**). The validation screen was performed on the CHLA06ATRT, D283, and UW228 cell lines, and data for each cell line were normalized to the 527 positive control pan-lethal gRNAs as described above. In the secondary screen, because the number of gRNAs for each gene varied, a scoring candidate was defined as a gene in which at least 30% of the gRNAs achieved a normalized abundance of less than or equal to −0.4. This threshold reflected the point that >95% of all negative control gRNAs failed to achieve in all 3 cell lines but >75% of all positive control gRNAs successfully achieved in all 3 cell lines. gRNAs were then grouped into their respective genomic region (e.g. UTR, ORF exon, adjacent gene exon, intron). Genes were then classified in the following manner according to the viability effect of the gRNAs: “selective uORF dependency” if only the ORF region gRNAs reached that threshold; “uORF and adjacent nucleotides” if the ORF gRNAs and gRNAs to only one other region of the RNA transcript scored; “uORF and CDS” if the ORF and an annotated adjacent protein coding gene both scored; “weak phenotype” if none of the cell lines showed a phenotype for that ORF.

### Base editing

gRNAs for base editing were manually designed to target the start codon of the uORF or associated parental CDS. The targeted nucleotide was positioned between basepairs 3 and 9 on the gRNA. gRNAs were synthesized via a commercial vendor (Synthego) with standard modifications (2’-O-Methyl at 3 first and last bases, 3′ phosphorothioate bonds between first 3 and last 2 bases). For base editing experiments, 200,000 D425 cells per reaction were centrifuged (1200RPM for 5 minutes), washed once in PBS, centrifuged again (1200 RPM for 5 minutes), and resuspended in 15uL of Nucleofector solution from the P3 kit (Lonza) in a 1.5mL microcentrifuge tube. Concurrent, a plasmid mix was prepared consisting of 1uL of Electroporation Enhancer (100uM, Lonza), 1.5 uL of 2ug/uL ABE8e-NRCH ribonucleoprotein editor,^83^ 1uL of base editor primer (50uM stock) and 3.6uL of Nucleofector supplement (Lonza). The ABE8e-NRCH base editor was a kind gift from Dr. David Liu’s lab at the Broad Institute. This 7.1uL of plasmid mix was added to the 15uL of cells in Nucleofector solution and samples were transferred to the Nucleocuvette vessels, ensuring that no bubbles were introduced in transfer. Cells were then electroporated using the Lonza DN-100 program. Afterwards, cells were recovered with the addition of 80uL of cell culture media. A cell count was repeated using a Beckman Coulter ViCell to ensure equal cell numbers and viability, and cells were transferred to 96 well poly-lysine coated plates at 2500 cells per well. Unused cells were plated on a 6 well poly-lysine coated plate and harvested for genomic DNA on day 4. Cell viability was measured at day 4 and day 6 using the Cell-Titer Glo assay (Promega). Viability data was analyzed by comparing the relative viability change between base editing with the uORF gRNA and the associated parental CDS gRNA. Negative controls were biological triplicate mock nucleofections.

The table below shows target sequences for base editing gRNAs, with PAM sites in italics, and the target start codon is in bold. The gRNA sequence is underlined.

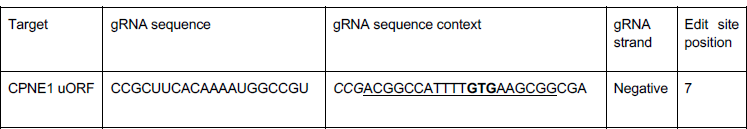
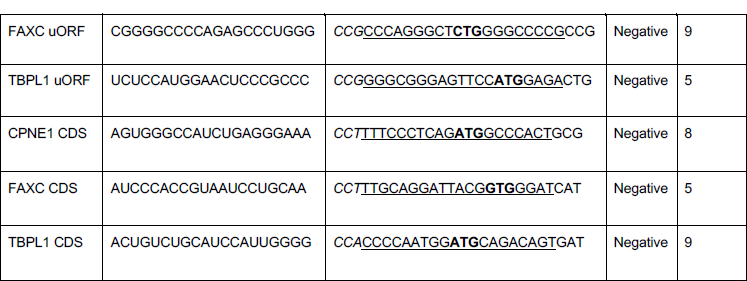

### Intergroup dependency comparisons in medulloblastoma and nomination of ASNSD1-uORF

To compare the overall impact for knockout of uORFs, uoORFs, and dORFs across molecular disease subtypes, the differential dependency for each ORF was assessed across each individual cell line. Individual values were averaged as the geometric mean across cell line subtypes as follows: MYC_medulloblastoma (D341, D283, D425, D458) and nonMYC (UW228, R256, ONS76) The distributions of differential dependency scores were compared across groups using a two-sided Student’s T test. For individual outlier uORFs, the weighted average of the differential dependency scores for uORFs and uoORFs for D283 and D341 were compared to those of UW228 and ONS76. Additionally, for each cell line, individual uORF outliers were assessed by calculating the delta differential dependency score between the uORF and the parental CDS and comparing this to the difference in the number of gRNAs that scored for the uORF compared to the parental CDS.

### ASNSD1-uORF evolutionary analysis

The amino acid sequence for ASNSD1-uORF (UniProt ID L0R819 isoform 1) and for the parental ASNSD1 CDS (UniProt ID Q9NWL6 isoform 1) were analyzed using the NCBI ProteinBlast feature (https://blast.ncbi.nlm.nih.gov/Blast.cgi?PAGE=Proteins) using default parameters against the “non-redundant protein sequences (nr)” database and the “model organisms (landmark)” database. All identified non-human amino acid sequences were downloaded and analyzed for similarity to either ASNSD1-uORF of ASNSD1 respectively using the ClustralOmega package (https://www.ebi.ac.uk/Tools/msa/clustalo/).

### ASNSD1 gene expression analysis

Processed RNA expression data for ASNSD1 mRNA expression (ENSG00000138381.9) were downloaded from GTeX for bulk RNA sequencing data (https://www.gtexportal.org/home/) and the Allen Institute Developing Brain Atlas (https://www.brainspan.org). In cell lines, ASNSD1 expression was evaluated through Cancer Cell Line Encyclopedia data for ASNSD1 (ENSG00000138381.9). CCLE data was downloaded from https://portals.broadinstitute.org/ccle. Data were analyzed in GraphPad Prism as shown.

### ASNSD1-uORF overexpression and rescue experiments

The indicated ASNSD1-uORF cDNAs were synthesized using a commercial vendor (GenScript) and cloned into the pLX_307 or pLX_313 mammalian expression vector (**Table S4M** for sequences). pLX_307 and pLX_313 are Gateway-compatible expression vectors where E1a is the promoter of the ORF and SV40 is the puromycin resistance gene with either puromycin (pLX_307) or hygromycin (pLX_313) resistance (details at https://portals.broadinstitute.org/gpp/public/resources/protocols). Lentivirus was produced in HEK293T cells as previously described, using the Lenti-X Concentrator (Takara Bio) to achieve a 50x virus concentration. For overexpression experiments, H9-derived NSC and D341 cells were transduced with lentivirus and stably-expressing cells were selected with either puromycin (0.5 ug/mL, plx_307 lentivirus) or hygromycin (300ug/mL, plx_313 lentivirus) for 72 hours prior to transitioning back to standard culture media. In 96 well plates (GelTrex pre-coated for H9-derived NSC or poly-lysine for D341), 4000-5000 cells per well were plated. For H9-derived NSC experiments, cell viability was monitored daily using the Cell-Titer Glo reagent. For D341 experiments, cells were infected with the indicated gRNA lentivirus 4-6 hours after plating. 16 hours after infection, cells were selected with 1ug/mL puromycin for 48 hours and grown for 7 days prior to cell viability analysis using CellTiter-Glo reagent.

### ASNSD1-uORF knockout experiments

Cells were plated in 96-well plates and allowed to grow for 4-8 hours prior to infection with the indicated sRNA or treatment condition. 1,000 - 5,000 cells per well were plated depending on the cell line. gRNAs were obtained from the Broad Institute Genomic Perturbation Platform (Broad Institute, Cambridge, MA, USA) or from direct synthesis into the BRDN0003 or BRDN0023 backbone via commercial vendor (GenScript, Piscataway, NJ). sgRNA sequences are listed below:

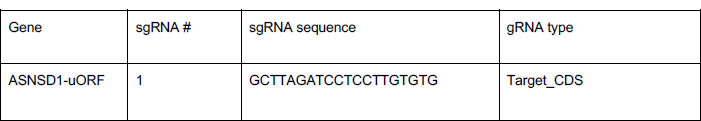
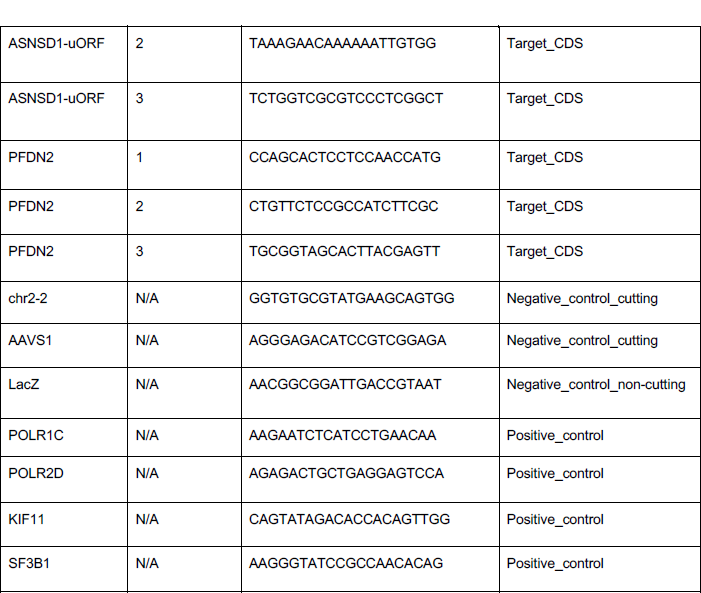

All sgRNAs were sequenced and verified. After sequence verification, constructs were transfected with packaging vectors into HEK-293T with Fugene HD (Sigma-Aldrich, St. Louis, MO). After plating, cells were then infected with sgRNA lentivirus to achieve maximal knockout but without viral toxicity. 16 hours after infection, cells were selected with 2ug/uL puromycin (Invitrogen, Carlsbad, CA) for 48 hours. Cell viability was measured CellTiter-Glo reagent (Promega, Madison, WI) was measured at 16 hours post-transfection for a baseline assessment, and additional timepoints as needed. For stable knockout cell lines, cells were plated at equal densities and cell viability was measured by CellTiter-Glo every 24 hours as indicated.

### Analysis of cell line knockout data

Cell line knockout data was normalized as previously described.^22^ Briefly, data for each cell line were standardized such that the average of the positive controls was equal to −1 and the average of the negative controls was equal to 0.

### Pooled ASNSD1-uORF knockout in the PRISM cell line panel

Pooled knockout screens in the PRISM cell line set were performed as previously described.^22^ Briefly, we used a pool of 486 barcoded human cancer cell lines, which were collectively grown in RPMI1640 media supplemented with 10% FBS. gRNAs used were non-cutting LacZ control (AACGGCGGATTGACCGTAAT), cutting control Chr2-2 (GGTGTGCGTATGAAGCAGTGG), ASNSD1-uORF #1 (GCTTAGATCCTCCTTGTGTG), and ASNSD1-uORF #2 (TAAAGAACAAAAAATTGTGG). Briefly, on Day 0, the cell pool was plated at 400,000 cells per well in a 6 well plate with a cell pellet collected for a “no infection” control. On Day 1, cells were transduced with gRNA and Cas9 using an all-in-one plasmid with lentiviral titer at an MOI of 10 and 4ug/mL polybrene. On Day 4, cell culture media was changed to include 1ug/mL puromycin for 72 hours, after which antibiotic-free media was used. Cells were then passaged every 72 hours and a cell pellet (2e6 cells) was collected for DNA on day 6, 10 and 15. For genomic DNA extraction, cell pellets were washed in PBS and then processed using the DNA Blood and Tissue Kit according to manufacturer’s instructions (Qiagen, Hilden, Germany).

For determination of individual cell line representation, DNA from each time point was amplified by PCR with universal barcode primers, and PCR products were confirmed on a 2% agarose gel for size. Then, PCR products were pooled and purified with AMPur beads (Beckman Coulter, Brea, CA), and DNA concentration was measured via Qubit fluorometric quantification (Thermo Fisher Scientific, Waltham, MA). DNA was sequenced on a NovaSeq (Illumina, San Diego, CA) at the Genomics Platform at the Broad Institute.

### Analysis of PRISM pooled ASNSD1-uORF knockout sequencing data

484 of 486 cell lines were detectable at the day 15 time point and were used for data analysis. Cell line abundance was determined by RNA expression of each cell line’s barcode using RNA-sequencing as previously described. Data analysis was performed as previously described^22^ with the following modifications: cell lines with a detected number of reads but with fewer than 12 reads were included in the analysis. Following calculation of reads, the log2 fold change abundance in each cell line was determined by comparing the day 15 abundance with the input plasmid pool. For lineage analysis of ASNSD1-uORF knockout across cancer types, we integrated the average log2 fold change of ASNSD1-uORF gRNA #1 and ASNSD1-uORF gRNA #2 with cancer cell line metadata from the DepMap database (www.depmap.org). For correlation of ASNSD1-uORF knockout phenotype with prefoldin complex knockout phenotypes, we used the Cancer Dependency Map release 21Q2 data to obtain gene-level knockout effects for 17643 human genes. A total of 389 cell lines were shared between the pooled ASNSD1-uORF knockout dataset and the Dependency Map dataset. For these 389 cell lines, the Pearson correlation was calculated for the knockout phenotypes relative to ASNSD1-uORF or members of the prefoldin and prefoldin-like complexes (PFDN1, PFDN2, PFDN4, PFDN5, PFDN6, URI1, UXT, PDRG1, VBP1) along with FDR-corrected Q values. The Pearson coefficients for each comparison were then permuted into a percentile rank and plotted as such. For evaluation of ASNSD1-uORF knockout with gene expression, the averaged ASNSD1-uORF knockout phenotype was compared to ASNSD1 mRNA expression (ENSG00000138381.9) using RNA-seq data values made available through the CCLE data at https://portals.broadinstitute.org/ccle.

### CRISPR-seq

The indicated cell lines were transduced with lentivirus for Ch2-2 or LacZ gRNA negative controls, ASNSD1-uORF gRNA #1 or ASNSD1-uORF gRNA #2. After selection of puromycin-resistant cells with 1 ug/mL puromycin for 48 hours, cells were grown until 96 hours post-transduction. Genomic DNA was then isolated from cells using the Qiagen DNeasy Blood & Tissue Kit (Qiagen, Hilden, Germany) according to the manufacturer’s instructors. 100ng of DNA was amplified by PCR with the following thermocycler conditions: 94C for 2 minutes, followed by 30 cycles of 94C for 30 seconds, 52C for 30 seconds, and 68C for 1 minute; final elongation was 68C for 7 minutes. PCR products were confirmed for specificity with a 1% agarose gel and then gel-purified with a Qiagen Gel Extraction kit according to manufacturer’s instructions. DNA was diluted to a concentration of 25ng/uL and submitted to the Massachusetts General Hospital Center for Computational and Integrative Biology (CCIB) DNA Core for sequencing. FASTQ sequencing files were analyzed using CRISPResso^84^ (http://crispresso.pinellolab.partners.org) according to default parameters.

Primers used for CRISPR-seq were:

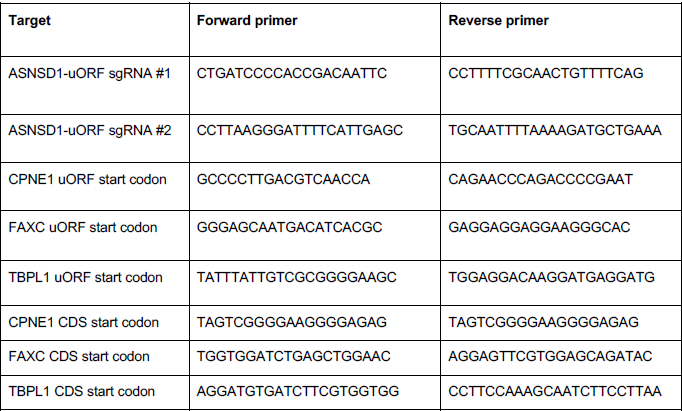

### ASNSD1 uORF protein abundance in cancer cell lines

Cancer cell lines were grown in standard tissue culture as previously described to a confluency of ∼80%. Cells were then washed three times in 1x ice-cold PBS, pelleted, and lysed using RIPA buffer. 35ug of cleared cell lysate was loaded per cell line on a 10-well, 10-20% Tris-Glycine gel and ran for 90 minutes at 125V. In each gel, samples were separated by an empty well. Then, the gels were washed 3x with deionized water at room temperature and stained with SimplyBlue Coomassie stain (Thermofisher) for 90 minutes at room temperature to ensure equal loading of protein. Gels were then washed 5 times with deionized water, 1 hour per wash at room temperature. Gel bands corresponding to the gel slice between 8 – 15 kDa were cut out using a sterile razor, started in 1mL of RNase/DNase free water, and then subjected to mass spectrometry analysis at the Taplin Mass Spectrometry facility at Harvard Medical School as previously described.^22^ Mass spectrometry data were normalized for individual proteins by calculating the fraction of that protein’s abundance relative to all proteins that were detected in that size range. The process was standardized using triplicate measurements for the D458 cell line. Additional cell lines were run in single replicates.

### ASNSD1-uORF abundance and correlations in mass spectrometry datasets

ASNSD1-uORF abundance was determined in publicly-available medulloblastoma mass spectrometry data^13^ as previously described.^22^ Data were obtained from the following repository: ftp://massive.ucsd.edu/MSV000082644. Briefly, a fasta file of the ASNSD1-uORF amino acid sequence was appended to the reference protein database. Raw mass spectrometry data were analyzed using Spectrum Mill v.7.09 (https://proteomics.broadinstitute.org). Search parameters, false discovery rate methodologies, and detailed descriptions for mass spectrometry datasets can be found in Prensner et al.^22^ Next, individual protein abundances were correlated to ASNSD1-uORF abundance using Pearson correlation coefficients and statistical significance of each correlation was corrected for multiple hypothesis testing by calculation of a q-value. Full values are available in **Table S4B**. For comparison of ASNSD1-uORF, PFDN1, PFDN2, PFDN4, PFDN5, PFDN6, VBP1, URI1, UXT, and PDRG1 abundance to MYC and MYCN levels, the maximum value of MYC or MYCN protein abundance was used, given their mutual exclusivity (Figure S4D). Then, samples were divided into quartiles based upon the maximum MYC or MYCN protein abundance for the 45 mass spectrometry samples, with N=11 samples in Quartiles 1, 2 and 3 and N=12 samples in Quartile 4. Data were normalized across the average of all samples to define the fold upregulation of Quartile 4 compared to all samples. Skew in protein levels was statistically determined using a 1-way ANOVA p value on GraphPad PRISM.

### Murine orthotopic xenograft experiments

Animal experiments were performed after approval by the Broad Institute and the Dana-Farber Institutional Care and Use Committee (IACUC) and were conducted as per NIH guidelines for animal welfare. Animals were housed and cared for according to standard guidelines with free access to water and food. All experiments were performed on 7 weeks-old female NSG mice (NOD.Cg-Prkdcscid Il2rgtm1Wjl/SzJ, The Jackson Laboratory, Bar Harbor, ME). Mice were euthanized as they developed neurological symptoms. To perform xenografting experiments, animals were injected intraperitoneally with the analgesic buprenorphine 0.05 mg/kg and then anesthetized with isoflurane 2– 3% mixed with medical air and placed on a stereotactic frame. Next, a small incision and a small burr hole was made with a 25-gauge needle and D425 cells (60,000 cells in 1 µL PBS) were injected stereotactically into the cerebellum (stereotactic coordinates zeroed on bregma: −1.0 mm X (ML), −7.0 mm Y (AP) and −2.5 mm Z (DV)) of 7 weeks-old female NSG mice at rate of 1 µL/min with use of an infusion pump before the incision was closed. Mice were then checked daily for signs of distress, including seizures, weight loss, or tremors, and euthanized as they developed neurological symptoms, including head tilt, seizures, sudden weight loss, loss of balance, and/or ataxia. Mouse brains collected at the survival endpoint were either fixed in 4% paraformaldehyde for 24 hours and subsequently stored in 70% ethanol and stored at room temperature, or snap-frozen on dry ice and stored at −80 °C.

### Murine magnetic resonance imaging

MRI was performed using a Bruker BioSpec 7T/30 cm USR horizontal bore Superconducting Magnet System (Bruker Corp.). This system provides a maximum gradient amplitude of 440 mT/m and slew rate of 3,440 T/m/s and uses a 23 mm ID birdcage volume radiofrequency (RF) coil for both RF excitation and receiving. Mice were anesthetized with 1.5% isoflurane mixed with 2 L/min air flow and positioned on the treatment table using the Bruker AutoPac with laser positioning. Body temperature of the mice was maintained at 37 °C using a warm air fan while on the treatment table, and respiration and body temperature were monitored and regulated using the SAII (Sa Instruments) monitoring and gating system, model 1025T. T2-weighted images of the brain were obtained using a fast spin echo (RARE) sequence with fat suppression. The following parameters were used for image acquisition: repetition time (TR) = 6,000 ms, echo time (TE) = 36 ms, field of view (FOV) = 19.2 x 19.2 mm2, matrix size = 192 x 192, spatial resolution = 100 x 100 µm2, slice thickness = 0.5 mm, number of slices = 29, rare factor = 16, number of averages = 8, and total acquisition time 7:30 min. Bruker Paravision 6.0.1 software was used for MRI data acquisition, and tumor volume was determined from MRI images processed using a semiautomatic segmentation analysis software (ClinicalVolumes).

### Murine *in utero* electroporation experiments

*In utero* electroporation (IUE) experiments were performed as previously described.^62, 63^ Briefly, mouse medulloblastomas are formed by the introduction of cDNAs expressing MYC and dominant negative p53 (DNp53). PiggyBac transposase DNA plasmids have luciferase and an IRES-GFP site for continuous GFP expression. We tested two conditions: DNp53 + MYC and DNp53 + MYC + ASNSD1-uORF. Both conditions included the pCAG-PBase transposase plasmid to stably integrate cDNA expression constructs. Specifically, 1 μg of concentrated DNA plasmid mixtures (1 μg/μL containing 0.05% Fast Green (Sigma)) was injected into the 4th ventricle of E13.5 mouse embryos using a pulled glass capillary pipette. Following DNA injection, embryos were electroporated by applying 5 pulses (45 V, 50 ms pulses with 950 ms intervals) with a 3 mm tweezer electrode positioned at the upper rhombic lip and cerebellar ventricular zone. Once born, pups were imaged via IVIS for luciferase at 1-2 weeks of age to identify successfully electroporated offspring. Mice were monitored every 3 days for new tumor-related neurologic symptoms (e.g. hydrocephalus, altered gait, lethargy, weight loss). Mice with symptoms were then euthanized according to IACUC guidelines. Tumor burden was be confirmed with GFP immunohistochemistry, using 50 uM tissue sections that are blocked in PBS + 0.5% Triton X-100 + 10% normal donkey serum prior to incubation with an antibody for eGFP (Aves, #GFP1020) and Hoechst (Thermo Fisher) for cell nuclei. 10 IUE tumor-bearing offspring were used per condition. The primary endpoint of time-to-death was analyzed using Kaplan-Meier curves with a log-rank test with a two-sided p<0.05 being significant. IUE experiments were performed under the University of Cincinnati IACUC approval protocol #16-07-06-01.

### ASNSD1-uORF immunoprecipitation

HEK293T cells were transiently transfected with ASNSD1-uORF-V5, ASNSD1-uORF-FLAG, ASNSD1-uORF deletion mutants (V5-tagged), GFP-V5 or GFP-FLAG fusion proteins using OptiMem and Fugene HD (Sigma-Aldrich). Forty-eight hours later, cells were washed once in ice-cold PBS and collected by centrifugation at 1,500 RPM for 5 minutes. Cells were lysed in lysis buffer (50 nM Tri-HCl pH 8.0, 150 nM NaCl, 2 mM EDTA pH 8.0, 0.2% NP-40 and 1 μg ml–1 PMSF protease inhibitor) for 20 minutes on ice and then cell debris was removed with centrifugation at 13,500 RPM for 15 minutes. Cell lysates were quantified using the BCA method and 2 mg of protein was used for input. Next, lysates were cleared with Pierce magnetic A/G beads (Thermo Fisher Scientific) for 1 h while rotating at 18–20 RPM. Beads were then discarded, and 10% of the medium was removed as an input sample and kept at 4 °C without freezing. The remaining culture medium was then treated with 50 μl of magnetic anti-V5 beads (MBL International) or 50uL of Anti-FLAG(R) M2 Magnetic Beads (Sigma-Aldrich) and rotated at 18–20 RPM overnight at 4°C. The following day, the supernatant was discarded and beads were washed four times in immunoprecipitation wash buffer (50 nM Tri-HCl pH 8.0, 150 nM NaCl, 2 mM EDTA pH 8.0, 0.02% NP-40 and 1 μg/ml PMSF protease inhibitor) with rotation for 10 min per wash. After the final wash, beads were gently centrifuged and residual wash buffer was removed. Then, proteins were eluted twice with 2 μg/μl V5 peptide in water (Sigma-Aldrich) or 1 μg/μl 3x FLAG peptide (ApexBio) at 37 °C for 15 min with shaking at 1,000 RPM The two elution fractions were pooled and samples were prepared with 4x LDS sample buffer and 10× sample-reducing agent (Thermo Fisher Scientific), followed by boiling at 95 °C for 5 min. One-third of the eluate was then run on a 10–20% Tris-glycine SDS–PAGE gel and stained with SimplyBlue Coomassie stain (Thermo Fisher Scientific) for 2 h. Gels were destained with a minimum of three washes in water for at least 2 h per wash. Bands were visualized using Coomassie autofluorescence on LI-COR Odyssey in the 800-nM channel. Gel lanes were then cut into six equal-sized pieces using a sterile razor under sterile conditions, and stored in 1 ml of RNase/DNAse-free water before LC-MS/MS analysis.

### PFDN6 co-immunoprecipitation and mass spectrometry

D425 medulloblastoma cells were grown to 80% confluency to ∼90 million cells. Cells were collected and washed twice in ice-cold PBS. Cells were lysed in endogenous IP lysis buffer (50 nM Tri-HCl pH 8.0, 150 nM NaCl, 2 mM EDTA pH 8.0, 0.2% NP-40, 2.5% Glycerol v/v 2.5%, Rnase I (1U/10uL) and Turbo DNase (25U/10uL), 1 μg/ml PMSF protease inhibitor). Lysis occurred for 15 minutes at room temperature and then 10 minutes on ice. Samples were centrifuged at 14000 RPM at 4C for 12 minutes to clear the lysates. Protein concentration was determined using the BCA method, and 200ug of input protein was saved for the input samples.. 2.5 mg of protein was aliquoted as the input for control IP and PFDN6 IP tubes, and samples were adjusted to 600uL with additional endogenous IP lysis buffer. Samples were mixed with 200uL of pre-washed slurry of a 1:1 mix of EZview Red Protein G and EZview Red Protein A bead affinity gel slurry (Sigma-Aldrich) and rotated at 4C for 1hr. Prior to usage, the protein A/G slurry was pre-washed 2x in endogenous IP lysis buffer. Samples were centrifuged at 250g x 4 minutes at 4C and supernatant was removed and kept in a new tube, with the beads discarded. This was performed twice to increase purity. Then, 20 uL of PFDN6 (Sigma-Aldrich #HPA043032) or normal rabbit IgG (Cell Signaling Technology #2729S) antibody was added to the appropriate tube, and samples were rotated at 18 RPM at 4C overnight. After overnight rotation, samples were incubated with 100uL EZview Red protein A/G bead slurry (1:1 mixture as above, pre-washed twice in IP wash buffer) for 2 hrs at 4C with 18 RPM Samples were centrifuged at 250g x 4 minutes at 4C and supernatant was removed, with the beads left behind. Beads were washed three times for 10 minutes each in ice-cold IP wash buffer with glycerol (50 nM Tri-HCl pH 8.0, 150 nM NaCl, 2 mM EDTA pH 8.0, 0.02% NP-40, 2.5% glycerol v/v and 1 μg/ml PMSF protease inhibitor). During each wash, samples were rotated at 18 RPM at 4C, and after each wash samples were centrifuged at 250g x 4 minutes at 4C and supernatant was removed. Samples were then eluted in 100uL of 1x sample loading buffer and boiled for 5 min at 95C. For mass spectrometry analysis, samples were run on a 16% Tris-Glycine gel at 125V for 100 minutes, then rinsed with deionized water and stained with SimplyBlue Coomassie stain (Thermo Fisher Scientific) for 2 hr. Gels were destained with a minimum of three washes in water for at least 2 h per wash. A gel slice corresponding to the band between 10 - 20 kDa was removed using a sterile razor under sterile conditions, and stored in 1 ml of RNase/DNAse-free water before LC-MS/MS analysis at the Taplin Mass Spectrometry Facility at Harvard Medical School. Experiments were performed in biological duplicate.

### Identification of downstream targets

1.5 million D425 cells or 2.0 million D283 cells were plated in each well of poly-lysine coated 6 well plates. Cells were allowed to attach for 3 hours and then subsequently transduced with 30uL of 10x concentrated lentivirus with 4ug/mL polybrene. Transductions were done in biological triplicate. Cells were grown for 24 hours and the 1.5ug/mL of puromycin was added. Cells were antibiotic-selected for 48 hours and then fresh media was added. Cells were grown for an additional 48 hours. At the 120 hour time point, cell media was aspirated and cells were washed in ice-cold PBS four times. Cells were scraped, counted, and aliquoted into 1 million cells for RNA-seq and 3 million cells for mass spectrometry. Cells were pelleted; PBS was removed and cells were flash-frozen in liquid nitrogen. RNA was isolated as above and mRNA sequencing was performed at the Dana-Farber Cancer Institute Molecular Biology Core Facility as above. RNA-seq read processing, alignment and quantification was performed as above. CDS read count normalization and differential expression analysis between knockout and control conditions was performed separately for each cell line using DESeq2. Cell pellets reserved for mass spectrometry were transferred to the Harvard Medical School ThermoFisher Center for Multiplexed Proteomics (TCMP) for total proteome analysis using TMT 10-plex or 15-plex. Protein lysates were subject to quantification, reduction and alkylation, precipitation and digestion followed by peptide quantification, TMT-labeling, LC-MS3 label check, basic reverse-phase HPLC fractionation (bRP-HPLC), LC-MS3 analysis of 12 bRP-LC peptide fractions, database searching, filtering to 1% FDR at protein level, TMT reporter quantification, and data analysis accord to standard TCMP core facility pipelines as previously described.^22^ To identify downstream targets, significantly differentially-abundant proteins with a p < 0.01 were considered. Proteins that had statistically-significant changes in both PFDN2 and ASNSD1-uORF knockouts were tested for gene network modules using the NCBI DAVID Bioinformatics platform (https://david.ncifcrf.gov/tools.jsp) on default settings.

### Prefoldin complex lethality in murine embryo knockout

Each subunit of the prefoldin and prefoldin-like complex was queried for mouse embryonic phenotypes using the information provided by the International Mouse Phenotyping Consortium.^85, 86^ Data were downloaded from https://www.mousephenotype.org and phenotypes observed in the homozygous knockout setting are reported.

### Comparison of CRISPR screen data with Project Achilles

The ASNSD1 gene was evaluated for cell line phenotypes using the DepMap_public_19Q4 release of CRISPR DepMap data and the Achilles RNA interference screens using the file “Achilles_logfold_change” (available at https://depmap.org/portal/download). Knockout phenotypes for 313 cell line assessed by both CRISPR and RNAi were z-scored and compared to each other.

### Statistical analyses for experimental studies

All data are expressed as means ± standard deviation. All experimental assays were performed in duplicate or triplicate. Statistical analysis was performed by a two-tailed Student’s t-test, one-way or two-way analysis of variance (ANOVA), Kolmogorov-Smirnov test, log-rank P value, or other tests as indicated. A p value <0.05 was considered statistically significant.

### Data availability

All raw sequencing data and custom code will be made publicly available upon publication. Upon publication, Ribo-seq and RNA-seq data for medulloblastoma cell lines, including RNA-seq following ASNSD1-uORF and PFDN2 knockout in D425 and D283 cells, will be available through the NCBI Short Read Archive through BioProject ID PRJNA957428. Ribo-seq and RNA-seq data for patient tissue samples from the Dana-Farber Cancer Institute are submitted to the NCBI dbGaP and will be made publicly available. Ribo-seq and RNA-seq data for patient tissue samples from the Princess Maxima Center are submitted to the European Genome-Phenome Archive (EGA) and will be made publicly available. Custom code for RNA-seq and Ribo-seq analyses is available through GitHub at https://github.com/damhof/hofman_et_al_2023_seq. Original western blots are available at Mendeley Data at https://data.mendeley.com/datasets/d63f7yzk3j/1.

## List of Supplementary Materials

### Supplementary Figures

Figure S1: Profiling RNA translation in medulloblastoma

Figure S2: Genomic perturbation of non-canonical ORFs in medulloblastoma to reveal ORF dependencies.

Figure S3: Characterization of ASNSD1-uORF as a genetic dependency in medulloblastoma.

Figure S4: Association of ASNSD1-uORF to the prefoldin-like complex in medulloblastoma.

### Supplementary Tables

Table S1: Details on medulloblastoma sequencing cohort.

Table S2: Results of non-canonical ORF CRISPR screens.

Table S3: Validation of ASNSD1-uORF in CRISPR assays.

Table S4: ASNSD1-uORF associates with the prefoldin-like complex in proteomic and genetic perturbation datasets.

---

**Figure S1.**
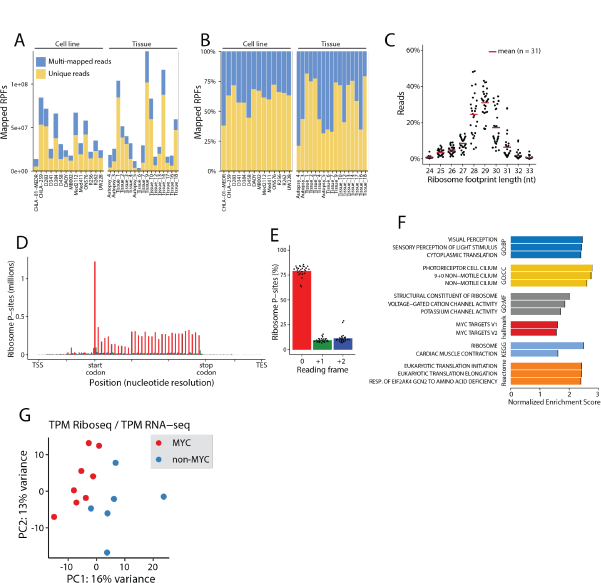
related to Figure 1: Profiling RNA translation in medulloblastoma. A) The absolute number of mapped ribosome protected footprints across each sample. Unique reads are indicated in yellow, and multi-mapping reads are indicated in blue. B) The relative proportion of unique and multi-mapping reads for each sample. C) Ribosome footprint sizes isolated for ribosome profiling. Each dot reflects a sample. The X axis shows the footprint size in nucleotides and the Y axis indicates the percentage of reads for each sample. D) A summarized plot of all ribosome profiling data, showing the in-frame P site periodicity across annotated protein-coding sequences. E) The average percentage of in-frame P-site reads, indicating >75% periodicity cumulatively for the dataset. F) Biological signatures enriched in genes that demonstrate differential translational efficiency in MYC-driven vs. non-MYC-driven medulloblastoma cell lines. G) A principal component analysis of the translational efficiency of non-canonical ORFs based on MYC-driven or non-MYC-driven status of medulloblastoma cell lines.

**Figure S2.**
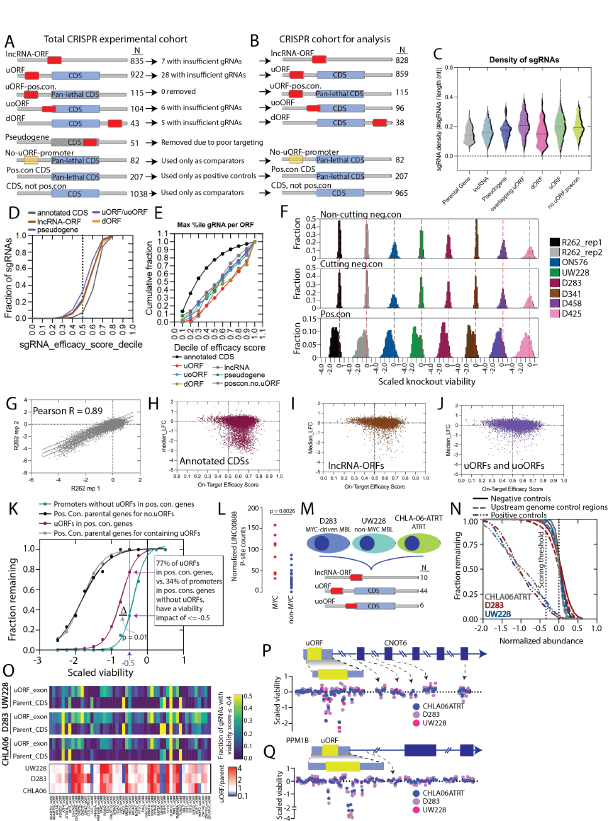
related to Figure 2: Genomic perturbation of non-canonical ORFs in medulloblastoma to reveal ORF dependencies. A) A schematic detailing all ORFs targeted by the custom CRISPR library and the number of cases which were removed from analysis. B) A schematic showing the final cohort of ORFs analyzed by CRISPR screening. C) A violin plot showing the density of gRNA design per length of ORF. D) The faction of gRNAs that achieve each decile of gRNA efficacy score. Results are displayed for each of the indicated ORF categories. E) The cumulative fraction of gRNAs targeting a given ORF biotype compared to the decile of the efficacy score for the gRNA with the least favorable characteristics. F) A histogram showing the scaled knockout viability effect for gRNAs targeting positive control CDSs (n = 1,654 gRNAs) compared to non-cutting gRNA controls (n = 503 gRNAs) or genome cutting gRNA controls (n = 497 gRNAs). Each cell line is shown in the indicated color. G) The correlation of gRNA knockout viability phenotypes for the two replicates of R262. A Pearson correlation is shown. H) A scatter plot for annotated CDSs, showing the gRNA on-target efficacy score compared to the median log2 fold change in gRNA representation at the Day 14 timepoint for all cell lines. I) A scatter plot for lncRNA-ORFs, showing the gRNA on-target efficacy score compared to the median log2 fold change in gRNA representation at the Day 14 timepoint for all cell lines. J) A scatter plot for uORFs and uoORFs, showing the gRNA on-target efficacy score compared to the median log2 fold change in gRNA representation at the Day 14 timepoint for all cell lines. K) An analysis of gRNAs targeting promoters of positive control genes without uORFs compared to gRNAs targeting uORFs found in positive control genes. The X axis is the scaled viability for the gRNAs and the Y axis is the fraction of gRNAs achieving that viability threshold. P value by a Kolmogorov-Smirnov test. L) The abundance of P-site counts for an ORF in LINC00888 across the medulloblastoma dataset. P value by a two-tailed Student’s t-test. M) A schematic representation of the cell lines and ORF types targeted in the secondary CRISPR screen for gRNA saturation. N) Verification of controls in the secondary tiling CRISPR screen, showing the fraction of positive control or negative control gRNAs (on the Y axis) achieving the indicated scaled viability threshold on the X axis. O) *Top*, a heatmap for each of the 3 cell lines tested in the secondary tiling screen showing the fraction of gRNAs with a viability score of <= −0.4 for each pair of a parental CDS and the matched uORF or uoORF. *Bottom*, a heatmap showing the fold change in fraction of gRNAs with a viability score of <=-0.4 for each of the three cell lines, calculated as (Fraction of uORF gRNAs with a viability score <=-0.4) / (Fraction of CDS gRNAs with a viability score <= −0.4). P) A graphical representation of the tiling CRISPR screen data for CNOT6. Each dot reflects a gRNA. gRNAs are colored according to each of the three cell lines. The Y axis reflected scaled viability for each gRNA knockout. The X axis reflects genomic position of the gRNA relative to the shown gene structure. Q) A graphical representation of the tiling CRISPR screen data for PPM1B. Each dot reflects a gRNA. gRNAs are colored according to each of the three cell lines. The Y axis reflected scaled viability for each gRNA knockout. The X axis reflects genomic position of the gRNA relative to the shown gene structure.

**Figure S3.**
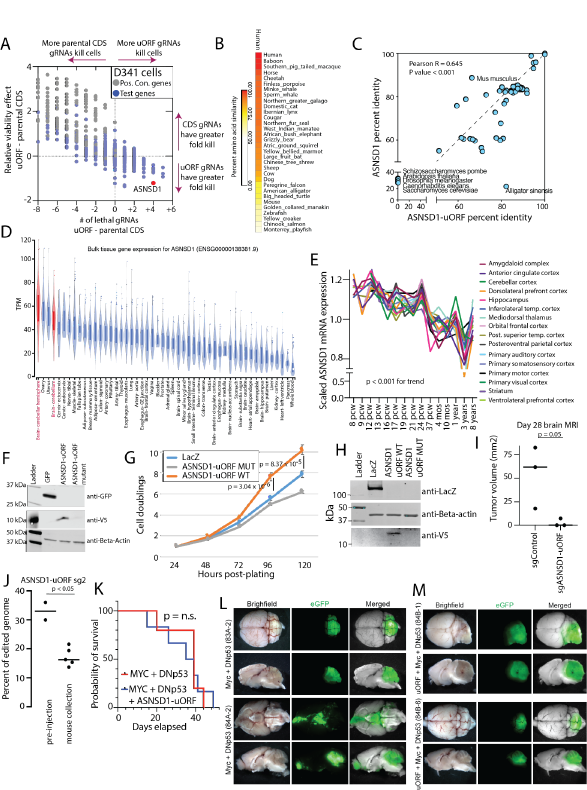
related to Figure 3: Characterization of ASNSD1-uORF as a genetic dependency in medulloblastoma. A) A scatter plot comparing the magnitude of viability phenotype of uORF knock-out relative to parental CDS knock-out in D341 cells. The X axis shows the number of gRNAs inducing a loss-of-viability phenotype for the uORF minus that number for the parental CDS. The Y axis shows the average loss-of-viability phenotype of the 4 most effective gRNAs for the uORF minus that number of the parental CDS. Positive control genes are shown in gray and other uORF genes are shown in blue. B) A heat map showing percent amino acid similarity between human ASNSD1-uORF and the amino acid sequences of its homolog in the indicated species. C) A scatter plot showing the percent amino acid similarity of homologs to ASNSD1-uORF to the human sequence (X axis) compared to the percent amino acid similarity of homologs of ASNSD1 to the human sequence (Y axis). Several species are highlighted if strongly discordant between the two proteins. D) ASNSD1 mRNA expression levels in the GTeX consortium. Cerebellar tissue is highlighted in red. Bulk tissue gene expression for ENSG00000138381.9 is shown. E) Normalized gene expression for ASNSD1 mRNA across human brain development. Data were obtained for ASNSD1 mRNA (ENSG00000138381.9) from the Allen Institute Developing Brain Atlas. P value by a two-sided ANOVA test. pcw, post-conception week; mos, months. F) Western blot analysis of overexpression of V5-tagged ASNSD1-uORF in D341 cells. G) The impact of ectopic expression of ASNSD1-uORF on cell growth in H9-derived neural stem cells. P values by a two-tailed Student’s T test. H) Western blot analysis of overexpression of V5-tagged ASNSD1-uORF in H9-derived NSC cells. I) Orthotopic xenograft tumor volume for D458 medulloblastoma cells on Day 22 after cerebellar injection. Tumor volume determined by MRI. P value by a two-tailed Student’s T-test. J) Assessment of ASNSD1-uORF knockout efficiency and persistence of knockout in D458 murine xenograft experiments. Knockout efficiency was determined by CRISPR sequencing of the gRNA cut site. P value by a Student’s T-test. K) Kaplan-Meier survival curves for in utero electroporation experiments testing mouse survival and medulloblastoma formation with cerebellar injection of cDNAs encoding MYC with a dominant-negative p53 (DNp53), either with or without addition injection of a cDNA encoding ASNSD1-uORF. P value by a log-rank test. n.s., non-significant. L) Whole brain images with GFP fluorescence for two mice with MYC and DNp53 induced medulloblastomas. M) Whole brain images with GFP fluorescence for two mice with MYC, DNp53 and ASNSD1-uORF medulloblastomas.

**Figure S4.**
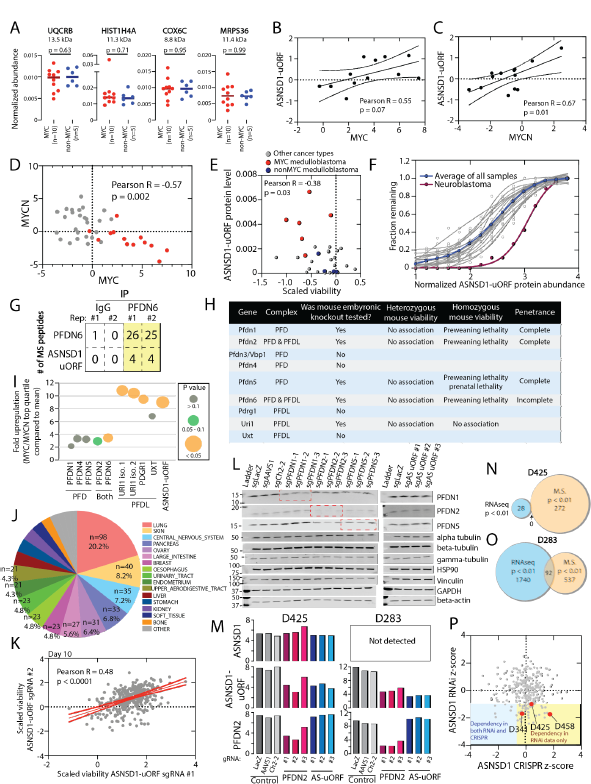
related to Figure 4: Association of ASNSD1-uORF to the prefoldin-like complex in medulloblastoma. A) Protein abundance of small protein controls in a cohort of MYC-driven (n=10) or non-MYC (n=5) medulloblastoma cell lines. P values by a two-tailed Student’s T test. B) Protein abundance of ASNSD1-uORF compared MYC in Group 3 medulloblastoma tissue samples only. Each dot reflects a sample. The Pearson correlation is shown. C) Protein abundance of ASNSD1-uORF compared MYC in Group 4 medulloblastoma tissue samples only. Each dot reflects a sample. The Pearson correlation is shown. D) RNA expression of MYC compared to MYCN in the medulloblastoma tissue cohort. Data obtained from^13^. Red dots reflect Group 3 medulloblastoma patients. A Pearson correlation is shown. E) A scatter plot showing the magnitude of viability loss after ASNSD1-uORF knockout (X axis) with the normalized ASNSD1-uORF protein abundance determined by mass spectrometry for 32 cell lines. Red dots are MYC-driven medulloblastoma cell lines. Blue dots are nonMYC medulloblastoma cell lines. Gray dots are non-medulloblastoma cell lines. F) Protein abundance of ASNSD1-uORF across the proteomics dataset for the Cancer Cell Line Encyclopedia (CCLE). The X axis shows the normalized protein abundance and the Y axis shows the fraction of samples within a given cell line lineage with the corresponding protein abundance. The red line indicates neuroblastoma cell lines, and gray lines reflect other cancer lineages. The blue line is the average of the dataset. G) Raw numbers of total peptides for co-immunoprecipitation experiments for endogenous PFDN6. Two replicates are shown. H) A table showing mouse germline knockout phenotypes for the indicated prefoldin or prefoldin-like complex proteins. I) Upregulation of prefoldin-like proteins in medulloblastoma tissue samples with high MYC/MYCN levels. The Y axis shows up regulation of the indicated protein compared to the mean. Size and color of the circles indicates degree of statistical significance of upregulation. P values by an ANOVA test. J) The representation of cancer cell line lineages across the 484 cell lines in the pooled knockout experiment. K) A scatter plot showing correlation of individual ASNSD1-uORF gRNAs used in the pooled knockout experiment. Each dot reflects a cell line. A Pearson correlation is shown for scaled viability values obtained on Day 10 after knockout. L) Western blot analysis of proteins reported to be regulated by the prefoldin complex following knockout of prefoldin proteins or ASNSD1-uORF in D425 medulloblastoma cells. M) Quantification of ASNSD1-uORF and PFDN2 protein knockout in D425 and D283 cells in shotgun mass spectrometry experiments. ASNSD1 parent CDS protein levels are also shown. N) The overlap and number of genes identified as regulated in mass spectrometry or RNAseq following ASNSD1-uORF knockout in D425 cells. The p values refer to the thresholds used to determine regulated genes in either mass spectrometry or RNAseq data. O) The overlap and number of genes identified as regulated in mass spectrometry or RNAseq following ASNSD1-uORF knockout in D283 cells. The p values refer to the thresholds used to determine regulated genes in either mass spectrometry or RNAseq data. P) A scatter plot showing loss-of-viability data for ASNSD1 in 313 cell lines tested by both RNA interference screening from Project Achilles^59^ and CRISPR in the cancer Dependency Map (www.depmap.org). Three medulloblastoma cell lines are highlighted.

